# Impact of N-terminal dimerization on formin homology 1 domain polymer dynamics and actin assembly

**DOI:** 10.1101/2025.11.21.689827

**Authors:** Katharine Bogue, Bryan Christian, Margot Quinlan, Jun Allard

**Affiliations:** Department of Chemistry & Biochemistry, University of California Los Angeles, Los Angeles, CA 90095-1569 USA; Department of Physics & Astronomy, Department of Mathematics, University of California Irvine, Irvine, CA 92697 USA

## Abstract

Many proteins contain intrinsically disordered regions (IDRs) that lack stable 3-dimensional structure. IDR behavior is poorly understood, leading to challenges for biochemical and computational analysis of IDR-containing proteins. Formins are a diverse set of homodimers containing an IDR — the FH1 domain — that facilitates polymerization of the cytoskeletal protein actin by increasing the local concentration of actin monomers at the actin assembly site. A commonly accepted model of formin-based actin polymerization involves a capture-and-deliver process: one or more binding sites (proline-rich motifs, PRMs) “capture” actin monomers and then “deliver” actin to the actin assembly site. There is evidence that formin FH1 domains are dimerized on both ends, but much research has been performed with formin constructs lacking the N-terminal dimerization site. Here, we ask: What happens when N-terminal dimerization is added to the standard model of formin-mediated actin assembly? We extend the kinetic model of FH1-mediated actin polymerization by incorporating a coarse-grain polymer model of FH1 domain dynamics, modeling the FH1 domain as a freely-jointed chain. We find that N-terminal dimerization can impact polymerization rates by modifying binding site accessibility and/or local concentration of binding sites (PRMs) at the actin assembly site (FH2 domain). Which effect dominates depends on kinetic parameters and formin properties such as FH1 domain length and binding site location. Additionally, we demonstrate that our model can be fit to experimental data and used to make predictions for the effects of N-terminal dimerization on a variety of formin family members.

**Significance:** Intrinsically disordered regions (IDRs) are common protein components that lack stable 3D structures and are thus difficult to study. Here, we develop a polymer-physics based computational model of the formin FH1 domain, an IDR involved in building the cell’s cytoskeleton. Some disease-associated mutations of formins occur in a region that is suspected to induce dimerization, forcing the FH1 domains to form a loop. Little is known about the loop’s prevalence, location, or impact on cytoskeletal assembly. Using simple polymer physics, we demonstrate that this dimerization can alter FH1 domain polymer dynamics and thus impact cytoskeletal assembly. These results not only highlight an important aspect of FH1-mediated cytoskeletal assembly, but also provide a framework for modeling IDRs.

## 1 Introduction

Intrinsically disordered regions (IDRs) are highly flexible stretches of protein characterized by a lack of stable structure. Roughly 44% of human proteins contain IDRs longer than 30 amino acids, and many IDR-containing proteins such as *ω*-Synuclein, BRCA-1, Hirudin, and Amylin have important disease implications (1, 2). The study of IDRs is challenging, in part because the traditional “structure-function” logic fails to describe such flexibility (1–3). Furthermore, the dynamic behavior of IDRs poses particular challenges for computational work, as the large number of possible conformations results in expensive simulations (1). Thus, a reliable computational method for characterizing individual IDR behavior is needed.

One example of an IDR-containing protein is provided by the formin family of proteins. Formins fall into nine classes, each of which may be represented by multiple isoforms (4). Humans have 15 formins that play distinct structural roles and are implicated in a range of diseases (5–8). Formins nucleate and polymerize actin, thus supporting actin-based cytoskeletal structures and cell motility (5, 6, 9–11). Formins contain two main structures relevant to actin assembly, the formin homology (FH) 1 and 2 domains, shown schematically in Figure 1. Two FH2 domains form a donut shape that stimulates actin nucleation and binds and tracks the fast growing ends (barbed ends) of actin filaments. The two FH1 domains, which are intrinsically disordered and highly variable between isoforms and species, aid in actin polymerization by increasing the local concentration of actin monomers (5, 6, 9, 12).

**Figure 1.**
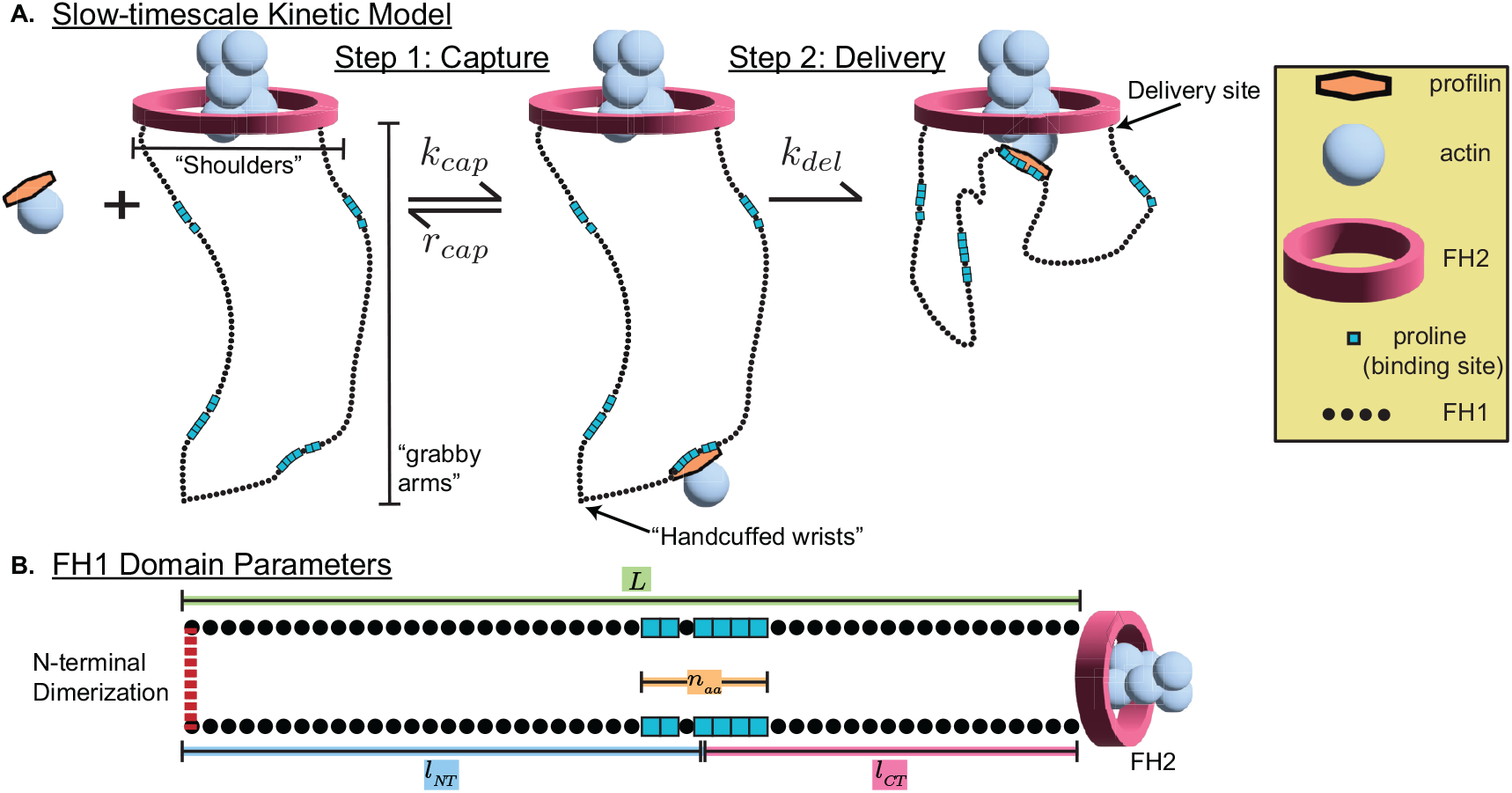
Model overview. (a) The 2 steps of the kinetic model represented schematically. In step 1, “capture” (*k*_cap_), binding sites (clusters of blue squares, PRMs, proline-rich motifs) along the FH1 domain (black circles) capture profilin-actin (orange + pale blue complex); this reaction is reversible (*r*_cap_). In step 2, “delivery” (*k*_del_), the FH1 domain delivers actin to the growing end of the actin filament, which is encircled by the FH2 domain (pink). The FH1-FH2 attachment point is considered to be the location of delivery, as noted with an arrow. (b) Model parameters are shown schematically: FH1 domain length (*L*, green), distance from binding site to FH2 domain (*l*_*CT*_, pink), distance from binding site to N-terminal dimerization site (*l*_*NT*_, blue), binding site size (*n*_*aa*_, orange), and N-terminal dimerization (dashed line). Note that only *n*_*aa*_ is a parameter in the kinetic model; All other parameters listed are relevant for the fast-timescale polymer simulations

Formin-mediated assembly of F-actin is well-described by a model in which each FH1 domain acts as a capture-and-deliver “grabby arm” (5, 6, 9–13). The FH1 domain contains proline-rich motifs (PRMs), which we will refer to as “binding sites” throughout the paper. The binding sites along the FH1 domain can bind to the actin-binding protein profilin. In the “capture” step, the FH1 domain “arms” reach out to capture profilin-actin complexes from solution. The FH1 domain arms then “deliver” these complexes to the growing actin filament at the FH2 domain “shoulders” (Figure 1).

A possible source of functional variation among formins may be the role of N-terminal dimerization (which can be visualized as “handcuffed wrists”). While all formins are dimerized at the C-terminal end by the FH2 domain, there is evidence that many formin family members are also dimerized at the remote end as well (5–7, 14). Many formins contain a dimerized Diaphanous inhibitory (DID) domain located in their N-terminal half (5–7, 14–16). Additionally, N-terminal coiled-coil domains downstream of the DID and closer to the FH1 domains are predicted in many formins (7). However, experimental validation of these coiled-coil domains is limited. We will refer to dimerization N-terminal to the FH1 simply as N-terminal dimerization from hereon. Notably, much experimental research uses C-terminal formin constructs lacking both the DID and the putative coiled-coil domain. Importantly, mutations in putative N-terminal dimerization domains are implicated in human diseases such as hypertrophic cardiomyopathy (17). The presence and relative location of N-terminal dimerization could have a significant impact on FH1 domain dynamics and thus FH1-mediated polymerization rates. One might predict that N-terminal dimerization would slow polymerization because the additional attachment point would increase steric hindrance of binding sites. However, if one considers the FH1 domains as entropic springs (18, 19), N-terminal dimerization doubles the entropic spring force pulling profilin-actin occupied binding sites to the FH2 domain, thereby accelerating polymerization. Alternatively, N-terminal dimerization may have negligible impact if, for example, the dimerization site is sufficiently far from the binding sites.

In this study, we ask: What happens when you consider the standard model of formin-mediated actin assembly in the context of N-terminal dimerization? To do so, we use computational simulation of the FH1 domain as an intrinsically disordered polymer. The model consists of a coarse-grain polymer model that explores FH1 domain configurations and a kinetic model simplified from Vavylonis et al. (12). The coarse-grain model predicts behavior of the FH1 domain on the fast timescale of microseconds to milliseconds (12, 13); The kinetic model predicts how capture and delivery give rise to overall polymerization rates on the slow timescale of milliseconds to seconds. Our model is agnostic to the secondary structure that produces N-terminal dimerization. Thus, the model allows for an understanding of the effects of dimerization due to a DID domain, coiled-coil, or other dimerization domain.

We find that there are 4 possible impacts of N-terminal dimerization (Figure 2 and Figure 3), depending on polymer behavior, kinetic parameters, and FH2 domain geometry: 1) deceleration due to decreased binding site accessibility, 2) acceleration due to increased local effective concentration of the binding site at the growing end, 3) acceleration due to increased binding site accessibility, and 4) deceleration due to decreased local effective concentration of the binding site at the growing end. Effect 4 dominates for realistic parameters. Lastly, we fit the model using experimental data. The budding yeast formin Bni1 was the first formin discovered to nucleate actin and is the most well-studied formin to date (6, 7, 20). The robust mechanistic information and experimental data available for Bni1 makes it an excellent formin to use for model fitting. Thus, we use experimental data acquired with a set of Bni1 constructs that contain a single binding site to fit the kinetic model (21). We then use these best-fit parameters to make predictions for the effects of N-terminal dimerization of a variety of formins.

**Figure 2.**
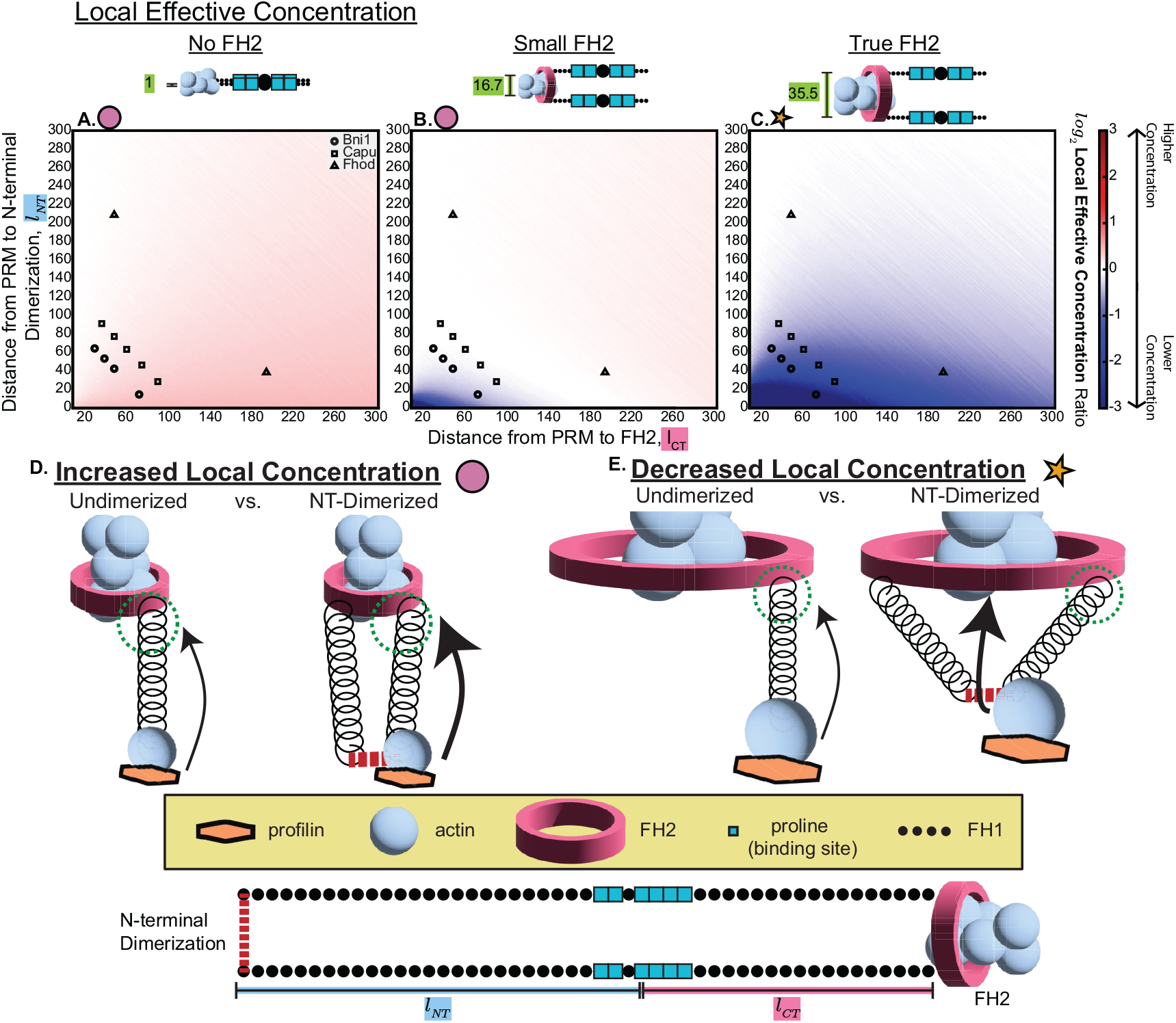
N-terminal dimerization differentially impacts FH1 domain local effective concentration. (a-c) Ratios (dimerized/non-dimerized) of simulated local effective concentration of binding sites at the growing end (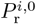) plotted as heatmaps with respect to the location of the binding site (PRM, proline-rich motif): distance from binding site to FH2 domain (*l*_*CT*_, x-axis) and distance from binding site to N-terminal dimerization site (*l*_*NT*_, y-axis). Higher (red) or lower (blue) local concentration of binding sites at the growing end enhances or impairs the rate of delivery, respectively. Simulations were run for 3 different FH2 domain sizes: (a) 1 amino acid, (b) 16.7 amino acids, and (c) 35.5 amino acids (from crystal structure of the Bni1 FH2 domain dimer, Xu et al. (22), PDB: 1UX5). Binding site locations for 3 formins are plotted over the heatmaps: Bni1 (circles), Capu (squares), and Fhod (triangles). (d-e) The two ways that N-terminal dimerization can impact polymerization via changes in local effective concentration are shown in schematics: (d) acceleration due to increased local effective concentration of the binding site at the growing end when the FH2 domain is small (pink circle, evident in a-b); (e) deceleration due to decreased local effective concentration of the binding site at the growing end when the FH2 domain is large (orange star, evident in c).

**Figure 3.**
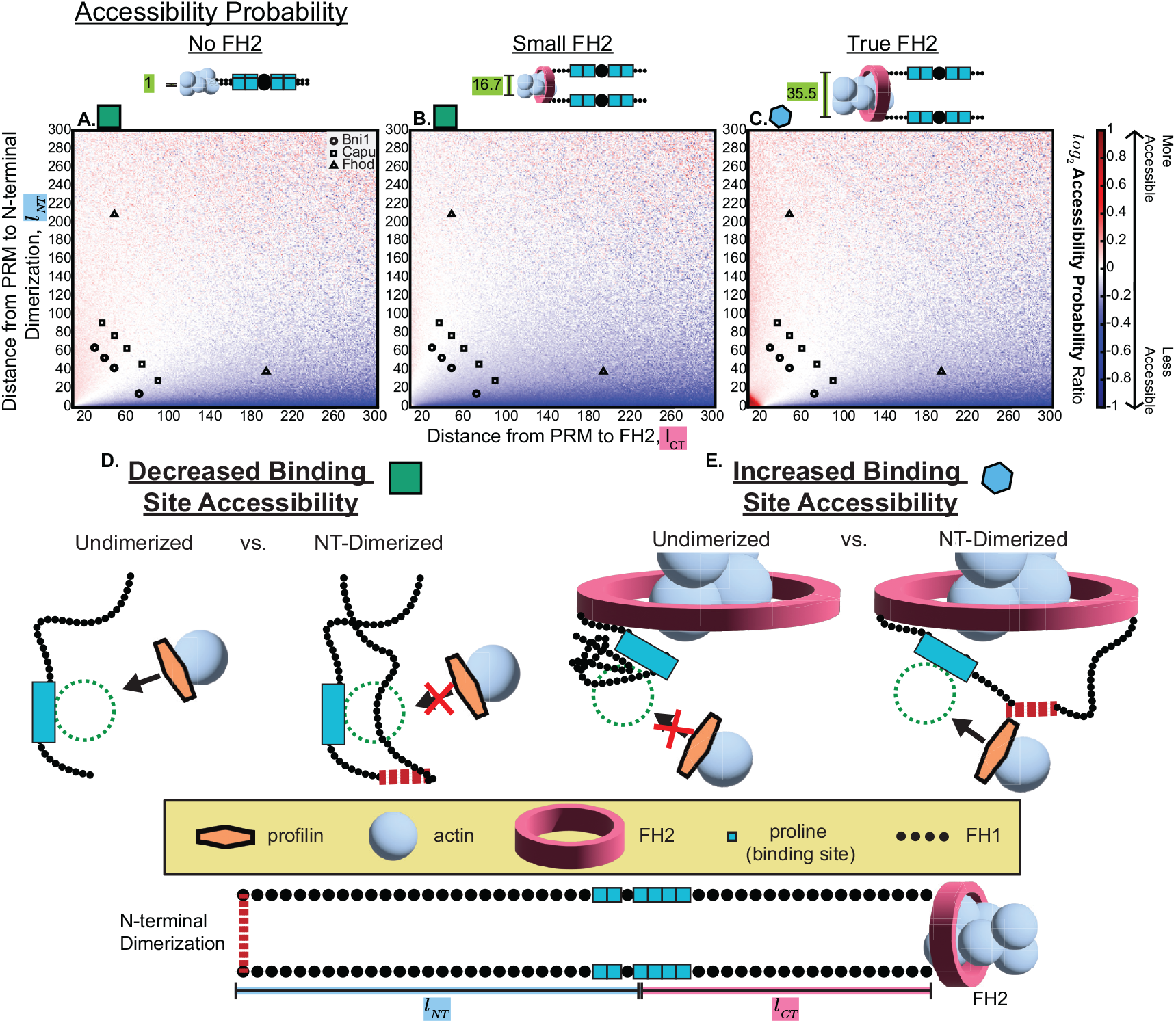
N-terminal dimerization differentially impacts FH1 domain accessibility probability. (a-c) Ratios (dimerized/non-dimerized) of simulated binding site probability of accessibility (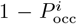) plotted as heatmaps with respect to the location of the binding site (PRM, proline-rich motif): distance from binding site to FH2 domain (*l*_*CT*_, x-axis) and distance from binding site to N-terminal dimerization site (*l*_*NT*_, y-axis). Increased (red) or decreased (blue) accessibility of binding sites enhances or impairs the rate of capture, respectively. Simulations were run for 3 different FH2 domain sizes: (a) 1 amino acid, (b) 16.7 amino acids, and (c) 35.5 amino acids (from crystal structure of the Bni1 FH2 domain dimer, Xu et al. (22), PDB: 1UX5). Binding site locations for 3 formins are plotted over the heatmaps: Bni1 (circles), Capu (squares), and Fhod (triangles). (d-e) The two ways that N-terminal dimerization can impact polymerization via changes in binding site accessibility are shown in schematics: (d) deceleration due to decreased binding site (PRM, proline-rich motif) accessibility when the FH2 domain is small (green square, evident in a-b), (e) acceleration due to increased binding site accessibility when the FH2 domain is large (blue hexagon, evident in c).

## 2 Methods

The dynamics of FH1-mediated actin polymerization occur on two timescales (12, 13), which we modeled separately and then combined. On the fast-timescale (microseconds to milliseconds), we modeled FH1 domain polymer dynamics using a coarse-grain polymer model. The coarse-grain model predicts parameters needed in the slow-timescale model: accessibility of binding sites and local effective concentration of binding sites at the assembly site. On the slow-timescale (milliseconds to seconds), we modeled profilin-actin-formin binding and unbinding using a kinetic model, which takes as input the fast-timescale behavior.

### 2.1 Coarse-Grain Polymer Model

Using the core polymer model described in Bryant et al. (13), we simulated the FH1 domain as a freely jointed chain of *N* rods with a Kuhn length of 1 amino acid (*L*_*k*_ = 3.6Å). Stationary anchor points of two identical chains were placed 35.5 Kuhn lengths apart (unless stated otherwise), representing the C-terminal end of each FH1 domain attached at the FH2 domain. The 35.5 Kuhn length distance was approximated from the distance between the N-terminal Lysine residues of a crystal structure of the Bni1 FH2 domain dimer (22) (PDB: 1UX5).

To explore the impact of FH2 domain size, chain base distances of 16.6 and 1 kuhn lengths were also used as noted. We performed two types of simulations: 1) “non-dimerized” simulations, where the N-terminal ends of the FH1 domains are free, and 2) “dimerized” simulations, where the N-terminal ends of the two FH1 domains are dimerized. N-terminal dimerization was imposed in the model by adding a dimerization domain with length of 1 Kuhn length.

We computed two equilibrium statistics of interest: 1) Binding site accessibility probability, 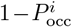, is the fraction of polymer configurations where no portions of the polymer are within one ligand radius from the binding site; 2) Probability density at the growing end, 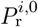, is the fraction of polymer configurations where the binding site is present within a specified radius of the delivery site, divided by the volume of that radius (see section 2.2 for a more detailed description of parameter notation). We used *r*_test_*/V*, where *r*_test_ is the test radius and 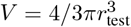 is the volume of the sphere of radius *r*_test_, such that the radius has units of µ M. We scale *r*_test_ linearly with *N*.

Equilibrium statistics for the tethered chains were determined using a Monte Carlo algorithm exploring the canonical ensemble (constant temperature). Each simulation was run until the distribution of root mean square end-to-end distance was converged with Kolmogorov-Smirnov statistic below 0.002, resulting in sample sizes ranging from about 10^7^ − 10^9^ conformations.

For dimerized simulations of long FH1 domains, *>* 200 − 300 amino acids (corresponding to the upper-right of each plot in Figure 2 and Figure 3), the Monte Carlo algorithm did not converge within our computational constraints. We found that the polymer behavior parameters of interest (binding site accessibility, 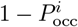, and local effective concentration at the growing end, 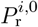) demonstrated smooth big-picture trends through FH1 domain lengths (*L*) of 600 amino acids despite lack of attaining the tight 0.002 threshold in the Kolmogorov-Smirnov statistic (see Figure 2 and Figure 3). However, out of an abundance of caution, we limited our predictions for individual formin polymerization rates (Figure 7) to formins with FH1 domain lengths (*L*) of 400 amino acids or less, which is sufficient to describe the majority of formin family members.

### 2.2 Kinetic Model

Equilibrium statistics from the coarse-grain polymer model were then used in the kinetic model to compute overall polymerization rates. The kinetic model is a simplified version of the Vavylonis et al. (12) model that consists of 3 states and 2 steps. Polymerization rates 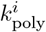 for individual binding sites (PRMs) are based on mean first passage time (MFPT; see appendix A and Figure 1). We use superscript *i* to refer to the ith binding site. We find that

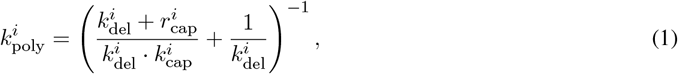

where *k*_cap_ is the rate of profilin-actin “capture” by a binding site, *k*_del_ refers to the rate of the profilin-actin occupied binding site “delivering” actin to the growing end of the growing actin filament, and *r*_cap_ refers to profilinactin falling off the binding site (see Figure 1).

The rate of capture for an individual binding site (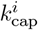) is modeled as

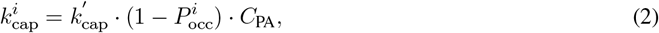

where 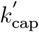 is the basal rate constant for capture, 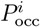 is the occlusion probability at the specified binding site (determined from the polymer simulations), and *C*_PA_ is the profilin-actin concentration in solution. The occlusion probability refers to the proportion of simulated polymer conformations in which the binding site is sterically occluded by other parts of the polymer, preventing profilin binding (13, 23, 24). We define the accessibility probability as the complement of occlusion probability (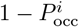). Since accessibility probability is proportional to polymerization rate, we use this term rather than referring to “1-occlusion probability” throughout the text. Additionally, we use the term “accessibility” as shorthand for “accessibility probability.”

The rate of delivery for an individual binding site (*k*^*i*^) is given by

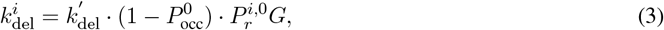

where 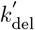 is the basal rate constant for delivery, 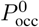 is the occlusion probability at the FH2 domain (determined from the polymer simulations), 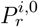 is the probability density of the binding site at the growing end (determined from the polymer simulations), and *G* is the formin-specific gating factor. The rate is proportional to (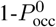), the probability that there is sufficient volume for the bound profilin-actin to attach at the delivery site (accessibility of the delivery site). The probability density, 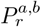refers to the probability of finding the binding site at a specific location. Here, the probability of finding a binding site at the growing end 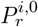 is used as a proxy for the local concentration of the binding site at the growing end (25–27). For clarity, we use the term “local effective concentration” to refer to probability density throughout the text.

The rate of reverse capture for an individual binding site (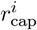) is the propensity that profilin-actin unbinds before successful delivery. Most of our simulations (Figure 2-Figure 5) use a constant 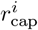. However, when fitting the model to data involving constructs with different binding sites (Figure 6) and when making predictions for formins with multiple binding sites (Figure 7), we assume 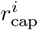 depends on binding site size as follows

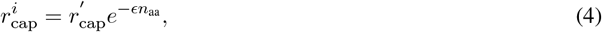

where 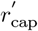 and ϵ are rate constants, and *n*_aa_ is the total number of amino acids in the binding site. In these cases,equation 4 assumes that the binding energy of profilin-actin is additive with each additional amino acid, which, by Detailed Balance, implies an exponential dependence of detachment rate.

The total rate of polymerization *k*_poly_ is computed as the sum of the polymerization rates of each binding site and of each arm of the FH1 domain homodimer. This additive assumption is valid as long as the delivery rates are below the capture rates, as appears to be the case (12, 13), despite some notable confounding reports (28, 29).

Heatmaps displaying overall polymerization rates — as determined by the kinetic model — in Figure 5 were smoothed using MATLAB’s Lowess Smoothing with a smoothing factor of 0.58.

### 2.3 Single Binding Site Bni1 Experiments

We fit the three-state model to data from experiments using the single binding site (PRM) Bni1 constructs described in Courtemanche and Pollard (21). We used WebPlotDigitizer (30) to approximate polymerization rates and error bar values for constructs with varied binding site locations (Figure 3 in Courtemanche and Pollard (21)) and binding site sizes (Figure 4 in Courtemanche and Pollard (21)). These experiments were performed with *Saccharomyces cerevisiae* profilin and 1.5µ M actin. We selected data from experiments performed at a profilin concentration of 5µ M, because these data represented the peak polymerization rates for the binding site size data and did not have any missing values. Using a binding affinity of 2.9µ M for *S. cerevisiae* profilin and actin, we calculated a corresponding profilin-actin concentration (*C*_PA_) of 0.88µ M for use in the kinetic model (31).

**Figure 4.**
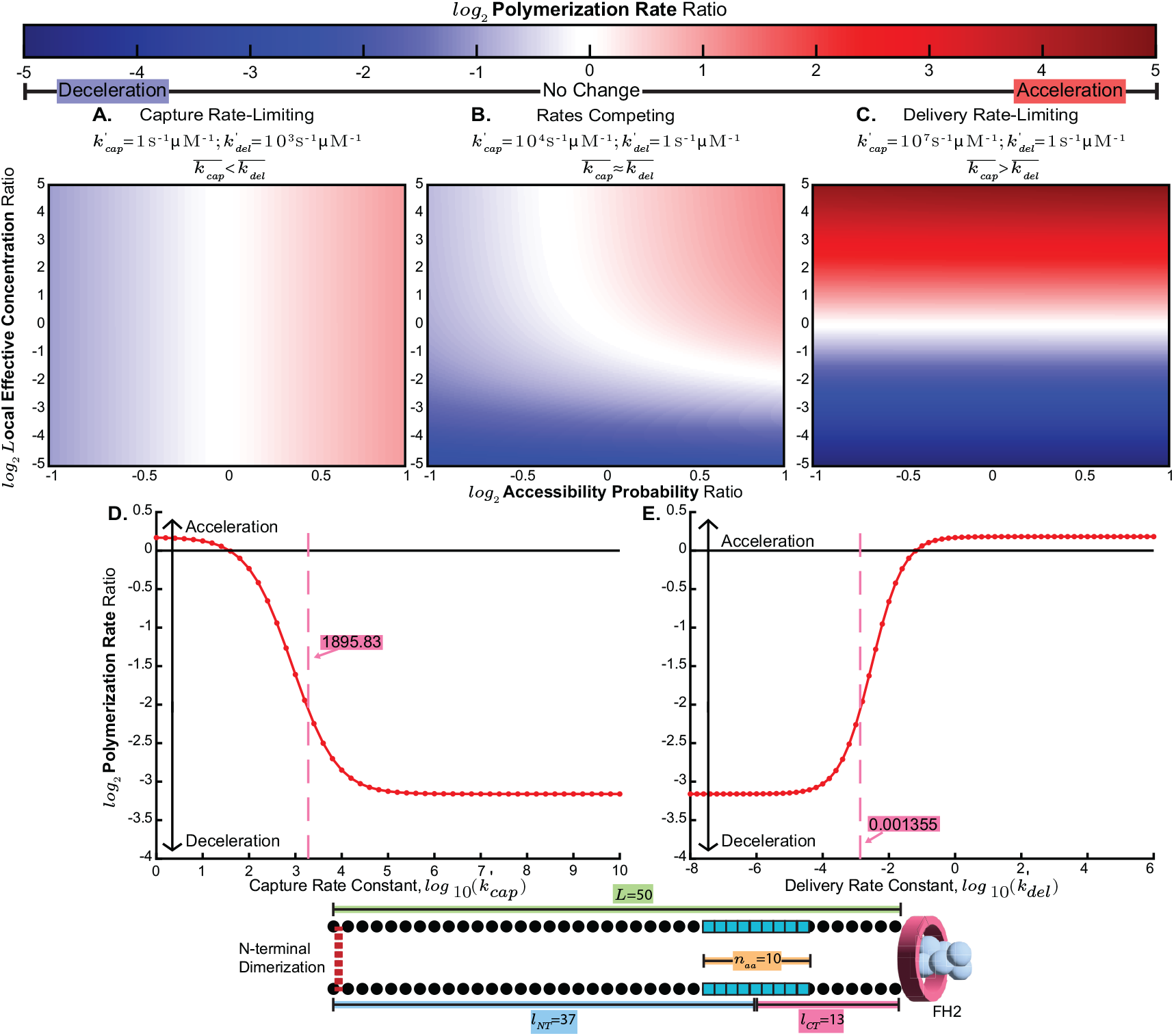
Changes in polymer behavior differentially impact polymerization rates based on the kinetic balance between capture and delivery rates. (a-c) Given the polymer behavior summarized in Figure 2 and Figure 3, the final polymerization rates depend on the kinetic rate parameters as illustrated by heatmaps of the ratio (dimerized/non-dimerized) of polymerization rates for a fictitious single binding site (PRM, proline-rich motif) formin. Heatmaps are plotted with respect to the ratios (dimerized/non-dimerized) of local effective concentration of binding sites at the growing end (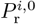, y-axis) and binding site accessibility (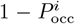, x-axis). Color-map values range from blue to red, indicating a deceleration or acceleration of polymerization due to N-terminal dimerization, respectively. Heatmaps were generated by fixing all other parameters and varying the local effective concentration and probability of accessibility for the dimerized formin. All regimes used the following parameters: 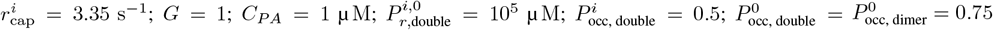. Values for 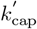 and 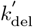 were selected to showcase the behavior when capture is rate-limiting 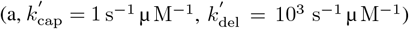, when delivery is rate-limiting 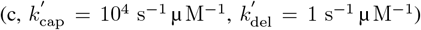, and when capture and delivery are competing (b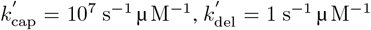). To clarify, these heatmaps show possible polymerization rates for arbitrary polymer behavior, as opposed to that which results from simulations. (d-e) Simulated polymerization rate ratios (dimerized/non-dimerized) of a single binding site fictitious formin construct with FH1 domain length *L* = 50, binding site size *n*_*aa*_ = 10, distance from binding site to N-terminal dimerization domain *l*_*NT*_ = 37, and distance from binding site to FH2 domain *l*_*CT*_ = 13. Ratios are plotted against changing 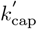 (d) and 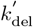 (e). Simulations were run using the following parameters (except when varied as indicated): 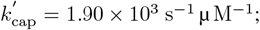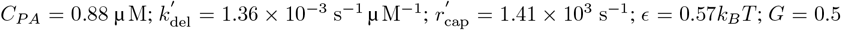. The locations of these best-fit parameters for each varied parameter are indicated by a dashed pink line.

### 2.4 Parameter Learning and Model Evaluation

In order to explore parameter space and identify best-fit parameters, we trained the model using experimental data. We performed Bayesian learning using Markov chain Monte Carlo (MCMC) with the Metropolis-Hastings algorithm (32–34) and likelihood function

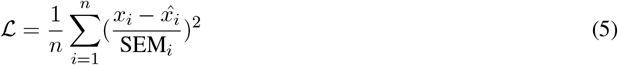

where SEM_*i*_ is the standard error of the mean, *n* is the number of data points, and *x*_*i*_ and 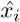 are the simulated and experimental data points, respectively. The Metropolis algorithm proposes random perturbations of fit parameters, accepting all new configurations that increase ℒ, and accepting some new configurations that decrease ℒ using a Boltzmann test (34). We used an adaptive step size that updates based on a target acceptance probability of 0.44 (34). We fit the model using data from the single binding site (PRM) Bni1 experiments (21). There were 4 fit parameters: 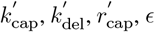. We bounded 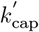 to be in the range of [10^−5^, 10^5^] s^−1^ µ M^−1^, 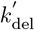 to be in the range of [10^−6^, 10^1^] s^−1^ µ M^−1^, 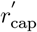 to be in the range of [10^2^, 10^20^] s^−1^ µ M^−1^, and ϵ to be in the range of [0.01, 4] *k*_*B*_*T*. We continued to generate proposals until the second third and last third of the accepted proposals for each fit parameter had approximately the same distribution with a Kolmogorov-Smirnov statistic *<* 0.02.

### 2.5 Formin and Formin Constructs

Formin FH1 domains were defined using UniProt (Table S1) or based on experimental constructs. The C-terminal residue of the FH1 domain was defined by the N-terminus of the FH2 domain as defined by UniProt or crystal structure (22). In our polymer dynamics simulations, we consider the FH1 domain’s N-terminal boundary to be the N-terminal dimerization domain. For sequences based on experimental Bni1 constructs (21), we defined the N-terminal residue of the FH1 domain as the N-terminal residue of the construct. For the sequences pulled from UniProt, we used a range of possible N-terminal dimerization domain locations (see Figure 6b-c and Figure 7) and thus a range of possible N-terminal residues of the FH1 domain.

### 2.6 Gating Factors

Gating factors for each formin were determined from existing literature. For gating factors used in predictions, see Table S1. For simulations of experimental Bni1 constructs, we used *G* = 0.5 (28).

### 2.7 Defining Binding Sites

Binding site (PRM) size and location were defined by scanning the FH1 domain sequence of a given formin from N to C termini and identifying instances of 4 or more sequential prolines with a maximum of 1 non-proline amino acid interruption (for example, this includes PPAPP, PPPP, and PAPPPP but not PAPP, PPP, or PPPAAP). We chose this PRM definition based on the demonstrated activity of the C-terminal PRM of Bni1 (pPD), which, with a sequence of PPAPP, is the smallest functional PRM of which we are aware (11, 21). We refer to the number of prolines in the binding site as *n*_*p*_ and the total length of the binding site, including any singular interruption, as *n*_*aa*_ (orange highlight in Figure 1b). The distance from the binding site to the FH2 domain, *l*_*CT*_, is the distance from the center amino acid in the binding site to the C-terminal amino acid in the FH1 domain (pink highlight in Figure 1b). The distance from the binding site to the N-terminal dimerization domain (*l*_*NT*_) is the distance from the N-terminal amino acid in the FH1 domain and the center amino acid in the binding site (blue highlight in Figure 1b). If the binding site has an even number of amino acids, the center amino acid is the N-terminal of the two center residues.

## 3 Results

The dynamics of FH1-mediated actin polymerization occur on two timescales (12, 13). To describe this, we developed a model for each timescale, and combined them. The fast-timescale model (microseconds to milliseconds) simulates FH1 domain polymer dynamics using a coarse-grain polymer model and predicts steric accessibility and local effective concentrations needed in the slow-timescale model. The slow-timescale model (milliseconds to seconds) simulates profilin-actin-formin binding and unbinding using a kinetic model and takes as input the fast-timescale behavior. After examining the behavior of each model separately, we combine the two models to yield a composite model to compute FH1-mediated polymerization rates.

### 3.1 Fast-timescale submodel: N-terminal dimerization influences FH1 local effective concentration and binding site accessibility

We used coarse-grain polymer simulations of FH1 domain as a freely jointed chain to explore how N-terminal dimerization impacts FH1 domain polymer behavior. Our polymer model predicts two key parameters that impact polymerization rates: 1) Binding site (PRM) probability of accessibility, 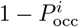, the probability that the binding site is not sterically occluded by other parts of the polymer, allowing for profilin binding; and 2) Probability density at the growing end, 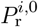, the local effective concentration of the binding site at the assembly site. These two parameters influence the rates of “capture” and “delivery,” respectively (see Figure 1 and Equation (2), Equation (3)). In order to assess the impact of N-terminal dimerization on accessibility and local effective concentration, we ran polymer simulations of N-terminally dimerized (hereafter, “dimerized”) and non-N-terminally dimerized (hereafter, “non-dimerized”) FH1 domains, connected by 3 possible FH2 domain sizes (“shoulder widths”). We ran simulations for FH1 domains of lengths (*L*) ranging from 1-600 amino acids and computed accessibility (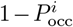) and local effective concentration (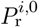) for binding sites located at all possible locations along the FH1 domain. The ratio of dimerized to non-dimerized values of local effective concentration (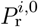) and accessibility (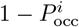) are shown as heatmaps with respect to binding site location in Figure 2 and Figure 3 respectively.

For a physiological FH2 domain size, we observed decreases in local effective concentration (Figure 2c) upon N-terminal dimerization. This outcome was unexpected, as, upon N-terminal dimerization, the entropic spring force pulling the FH1 domain towards the assembly site doubles, which we assumed would result in an increase in local effective concentration (Figure 2d). To better understand this phenomenon, we explored smaller, counter-factual FH2 domain sizes. Note that these counter-factual FH2 domain widths are used solely to aid our interpretation of simulations using a physiological FH2 domain size and are not intended to represent any real formins or biological consequences. Compared to simulations using the physiological FH2 domain size, simulations using the smallest FH2 domain sizes resulted in increases in local effective concentration (Figure 2a-b). Notably, the occurrence of decreased local effective concentration (size of the blue regions in Figure 2a-c) increases as FH2 domain size increases. Thus, the decrease in local effective concentration observed for physiological FH2 domain size must be explained by the size of the FH2 domain. Prior to N-terminal dimerization, the entropic spring force pulled each FH1 domain towards its respective delivery site (Figure 2d), which, following cryo-EM structure of the formin Cdc12, is “off-center” (35), i.e., not in the center of the barbed end. However, upon N-terminal dimerization, each FH1 domain is pulled towards the center. If the FH2 domain is small enough, this results in the expected increase in FH1 domain density at the delivery site (Figure 2d). However, when FH2 domain width is large, the anchors are moved farther away from the center of the FH2 domain. If the FH2 domain is large enough, this results in N-terminal dimerization pulling the FH1 domain away from the off-center delivery sites and decreasing local effective concentration (Figure 2e).

For a physiological FH2 domain size, we predominantly observed decreases (blue region of Figure 3c) or minimal/no change (light red and light blue “static” in the majority of Figure 3c) in accessibility upon N-terminal dimerization. Decreased accessibility was expected, as N-terminal dimerization brings the two FH1 domains in closer proximity, increasing the likelihood of portions of the FH1 polymer occluding binding sites (Figure 3d). However, we also observed increased accessibility for small FH1 domain lengths (red “blip” in the bottom left corner of Figure 3c). As with the unexpected behavior observed for local effective concentration, this unexpected behavior can be explained by exploring smaller, counter-factual FH2 domain sizes. Compared to simulations using the physiological FH2 domain size, simulations using the 1 and 16.7 residue wide FH2 domains do not produce the same increase in accessibility (Figure 3a-b). Moreover, the occurrences of increased accessibility (size of the red regions in Figure 3a-c) increase as FH2 domain size increases: At the largest FH2 domain size, we observed the most increase in accessibility (red “blip” in the bottom left corner of Figure 3c); This “blip” exists for the intermediate (16.7 amino acid) FH2 domain size but is almost imperceptible (Figure 3b), and is entirely absent for the smallest (1 amino acid) FH2 domain size (Figure 3a). Thus, the increase in accessibility observed for physiological FH2 domain size is also explained by the size of the FH2 domain. As previously discussed, as the FH2 domain size increases, FH1 domain density is pulled farther away from each FH1’s attachment point upon N-terminal dimerization. When the FH1 domain is short enough, this pull away from the attachment point results in a spreading out of the FH1 domain density, thus increasing binding site accessibility (Figure 3e).

In Figure 2 and Figure 3, FH1 length increases along the diagonal from the origin to the top left corner. The model predicted differences in accessibility (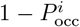) and local effective concentration (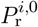) based on binding site location and FH1 domain length. For 16.7 and 35.5 residue wide FH2 domains, local effective concentration considerably decreases as FH1 domain length decreases (the blue region in the bottom left corner of Figure 2b-c deepens as distance from the origin decreases). We hypothesized that this behavior could be explained by the restriction in possible conformations upon N-terminal dimerization, due to the large size of FH2. In the extreme case, where the FH1 domains are shorter than or as long as the FH2 domain “shoulder width”, binding sites are likely unable to physically reach the delivery site after N-terminal dimerization. If true, this effect would only be present for finite FH2 domains. Consistent with our hypothesis, short FH1 domains do not display decreases in local effective concentration when the FH2 domain is very small (Figure 2a). Additionally, accessibility tends to decrease for binding sites near the N-terminal dimerization domain (bottom of Figure 3a-c) but experiences no change — or a very mild increase — for binding sites near the FH2 domain (left of Figure 3a-c). This is reasonable given how N-terminal dimerization brings the FH1 domains in closer proximity. Binding sites near the N-terminal dimerization domain are placed the closest to the other FH1 domain, and thus experience the strongest decrease in accessibility. This proximity effect diminishes as the binding site gets farther from the point of dimerization.

### 3.2 Slow-timescale submodel: The balance between the base rates of capture and delivery determines how polymer behavior impacts the effect of N-terminal dimerization

The above model describes how FH1 domain polymer properties influence local effective concentration and volume exclusion, and how these change due to N-terminal dimerization. To understand how these polymer properties impact the actin assembly rate, we used a simplified version of the formin model developed in Vavylonis et al. (Figure 1). Overall polymerization rates, *k*_poly_, are determined by the rates of capture, reverse capture, and delivery (Equation (1)). The rate of each of these steps is modulated by a rate constant, and the rates of capture and delivery include probability of accessibility (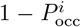) and local effective concentration (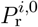), respectively (Equations (2) to (4)). In this section, we seek to understand the influence of N-terminal dimerization through the behavior of the kinetic model.

In order to study the impact of the kinetic parameter regime, we varied the rate constants for capture (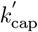) and delivery 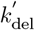 to produce three parameter regimes: capture rate-limiting (Figure 4a), delivery rate-limiting (Figure 4c), and a model where their rates are approximately equal (Figure 4b). We then varied the probability of accessibility and local effective concentration to establish the kinetic model’s dependence on each of these parameters (Figure 4a-c). We computed the polymerization rate ratio (dimerized/non-dimerized) for input probability of accessibility ratios ranging from a 50% decrease to a 100% increase in accessibility (−1 to 1 on a *log*_2_ scale) and input local effective concentration ratios ranging from −5 to 5 on a *log*_2_ scale. The behavior varies based on the overall parameter regime of the model: In parameter regimes where capture is the rate-limiting step (Figure 4a), the impact of N-terminal dimerization is dictated by binding site accessibility, with decreases in accessibility resulting in decreases in polymerization rates; In parameter regimes where delivery is the rate-limiting step (Figure 4c), the impact of N-terminal dimerization is dictated by the local effective concentration, with increases in local effective concentration resulting in increases in polymerization rates; In parameter regimes where capture and delivery compete with one another (Figure 4b), both binding site accessibility and local effective concentration affect changes in polymerization rates due to N-terminal dimerization. In the latter case, the changes in accessibility and local effective concentration determine whether capture or delivery is the rate-limiting factor: when the local effective concentration ratio is very large (N-terminal dimerization increases local effective concentration), capture dictates the effect of N-terminal dimerization and, when the probability of accessibility ratio is very large (N-terminal dimerization increases accessibility), delivery dictates the effect of N-terminal dimerization. In parameter regimes where capture and delivery compete with one another, acceleration-promoting polymer behavior in one parameter is dampened by deceleration-promoting polymer behavior in the other. Thus, increases in both capture and delivery are required for the overall polymerization rate to accelerate (light red top right corner of Figure 4b).

To further illustrate the impact of the balance between the capture and delivery rates, we computed the polymerization rate ratio for a fictitious formin across possible capture (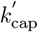) and delivery 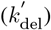 rate constants (Figure 4d-e). We chose a short (*L* = 50 amino acids) formin with a single large (*n*_*aa*_ = 10) binding site located near the assembly site (*l*_*NT*_ = 37 and *l*_*CT*_ = 13). Inputting these parameters into the kinetic model, we observed a sensitive parameter regime between the capture-rate-limiting and delivery-rate-limiting parameter regimes: when capture is slow or delivery is fast 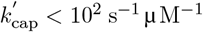 Figure 4d and 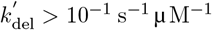 Figure 4e), N-terminal dimerization results in a constant mild acceleration; when capture is fast or delivery is slow 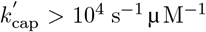 Figure 4d and 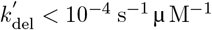 Figure 4e), N-terminal dimerization results in a constant moderate deceleration; when capture or delivery are of modest speeds (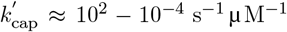 in Figure 4d or 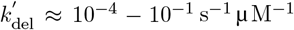), the effects of N-terminal dimerization rapidly change with the rate constants. Notably, the best-fit parameters from model fitting (described in section 3.4) fall in the sensitive regime where capture and delivery compete with one another (pink dashed lines Figure 4d-e). Overall, the model’s response to different balances between capture and delivery illustrate the driving forces behind acceleration and deceleration: Since acceleration occurs when capture is rate-limiting, increased capture rates (due to increased binding site accessibility) drive acceleration of overall polymerization; Since deceleration occurs when delivery is rate-limiting, decreased delivery rates (due to decreased local effective concentration) drive deceleration of overall polymerization.

### 3.3 Multiscale model: The overall influence of N-terminal dimerization on polymerization is complex but understandable through its parts

By combining the fast-timescale and slow-timescale models, we predicted how the FH1-mediated polymerization rate depends on FH1 domain polymer properties. In this composite model, we used the computed binding site accessibility (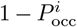) and local effective concentration (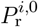) from our polymer simulations above (Figure 2 and Figure 3) as inputs to our kinetic model (Figure 4). The outcomes for 3 possible parameter regimes — capture rate-limiting, capture and delivery competing, and delivery rate-limiting — are shown in Figure 5. We plotted the ratio of dimerized to non-dimerized polymerization rates (*k*_poly_) with respect to binding site location and using the physiological FH2 domain width (35.5 residues). Results for counter-factual small FH2 domain size are shown in Figure S1. In all cases, both acceleration and deceleration of FH1-mediated polymerization are predicted depending on binding site location and parameter regime (Figure 5).

**Figure 5.**
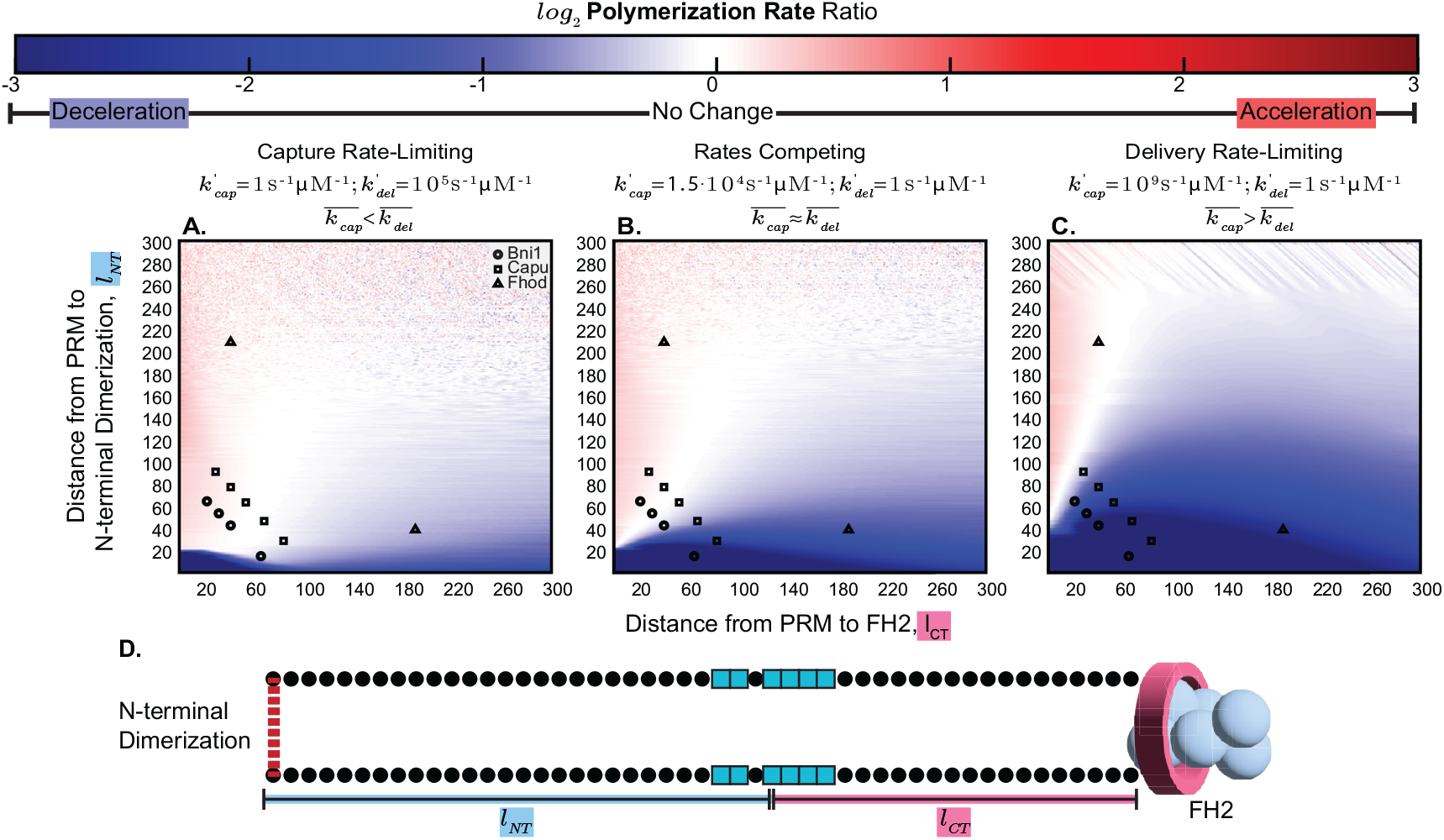
N-terminal dimerization can result in acceleration or deceleration of polymerization based on parameter regime. Combining the polymer behavior model (Figure 2 and Figure 3) with the kinetic model (Figure 4), we can predict the impact of N-terminal dimerization on overall polymerization rates. The ratio (dimerized/non-dimerized) of polymerization rates for formins with a single binding site (PRM, proline-rich motif) are plotted as heatmaps with respect to the location of the binding site: distance from binding site to FH2 domain (*l*_*CT*_, x-axis) and distance from binding site to N-terminal dimerization site (*l*_*NT*_, y-axis). Color-map values range from blue to red, indicating a deceleration or acceleration of polymerization due to N-terminal dimerization, respectively. All regimes used the following parameters: 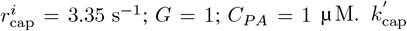 and 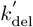 values were selected to showcase the behavior when capture is rate-limiting (a; 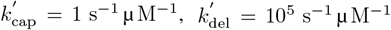), when delivery is rate-limiting (c; 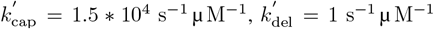), and when capture and delivery are competing (b 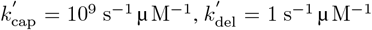). FH2 domain size of 35.5 amino acids was used (from crystal structure of the Bni1 FH2 domain dimer, Xu et al. (22), PDB: 1UX5). Binding site locations for 3 formins are plotted over the heatmaps: Bni1 (circles), Capu (squares), and Fhod (triangles).

The overall behavior that emerges from this simple model can be understood by breaking it down into its components. In all parameter regimes, N-terminal dimerization results in deceleration for binding sites close to the N-terminal dimerization domain (bottom of plots in Figure 5). This follows from the decrease in local effective concentration (bottom of Figure 2c) and decrease in accessibility (bottom of Figure 3c) for these binding sites. The region of the heatmaps demonstrating deceleration expands as delivery becomes a greater determinant of the overall polymerization rate. When capture is rate-limiting, the polymerization ratio is determined by changes in accessibility (Figure 4a), which decreases for binding sites near the N-terminal dimerization domain (bottom of Figure 3c) but increases for binding sites near the FH2 domain (left of Figure 3c). When delivery is rate-limiting, the polymerization ratio is determined by changes in local effective concentration (Figure 4c), which decreases for most binding site locations (Figure 2c). Thus, when capture is rate-limiting, the polymerization rate ratio heatmaps (Figure 5a) look more like the heatmaps for probability of accessibility ratio (Figure 3c), and, when delivery is rate-limiting, the polymerization rate ratio heatmaps (Figure 5c) look more like the heatmaps for local effective concentration ratio (Figure 2c). When capture and delivery compete with one another, the polymerization rate ratio heatmaps display a combination of these behaviors (Figure 5b). Interestingly, the strong increase in binding site accessibility observed for short FH1 domains (bottom left corner of Figure 3c) does not translate to an acceleration of polymerization when capture is rate-limiting (Figure 5a). This is because, although capture is generally rate-limiting in that regime, the magnitude of local effective concentration when the FH1 domain is short (bottom left corner of Figure 2c) is extremely decreased in comparison, thus making delivery the determining factor for these binding site locations. Predicted accelerations only occur when binding site accessibility increases and local effective concentration experiences a lesser change. These conditions appear to only occur when the increase in binding site accessibility is mild, resulting in only weak acceleration.

### 3.4 The model can be trained on experimental data in non-N-terminally dimerized Bni1 and used to predict the impacts of N-terminal dimerization

We trained the model with data from experiments using single binding site Bni1 constructs in Courtemanche and Pollard (21). We used a Bayesian learning algorithm (see section 2.4) to determine kinetic rate constants (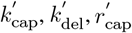) and binding site affinity (ϵ). Instead of returning a single best fit parameter set, Bayesian learning produces an ensemble of parameter sets, termed the posterior distribution, and a likelihood associated with each parameter set (32–34). Using the Bni1 data, the posterior distribution showed multiple regions of high likelihood (Figure 6a). This suggests that there are multiple parameter sets that fit the currently available data, from the perspective of likelihood.

**Figure 6.**
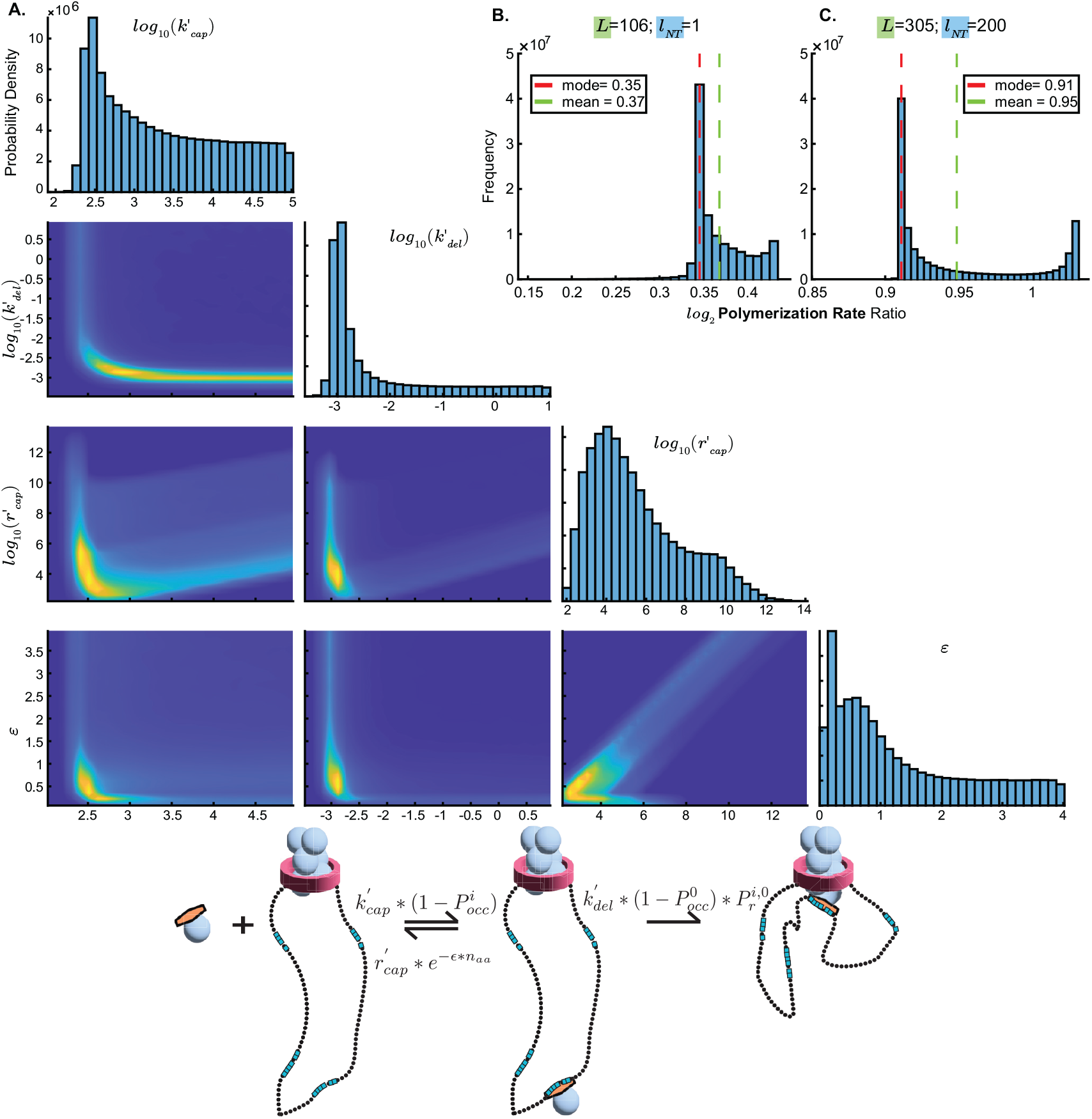
The model can be fit to single binding site Bni1 data and used to predict the impact of N-terminal dimerization. (a) Parameters learned by fitting the model to single binding site (PRM, proline-rich motif) Bni1 experimental data, shown as a posterior distribution. Highly likely parameter sets are shown in yellow, with parameter sets becoming less likely as the colors change from green, light blue, and dark blue. (b-c) Predicted polymerization rate ratios (dimerized/non-dimerized) for Bni1 using parameters from the posterior distribution of the MCMC fit to single binding site Bni1 data. Simulations were run for 2 different binding site locations (*l*_*NT*_) and FH1 domain lengths (*L*): (b) *L* = 106, *l*_*NT*_ = 1 and (c) *L* = 305, *l*_*NT*_ = 200. The histogram mode (red dashed line) and mean (green dashed line) are plotted on the histograms. For all simulations using Bni1 (or a Bni1 construct), the following parameters were used: *G* = 0.5; *C*_*PA*_ = 0.88 µ M.

The posteriors predicted both deceleration and mild acceleration of the polymerization rate of Bni1 upon N-terminal dimerization (Figure 6b-c). Since the location of the N-terminal dimerization domain of Bni1 is unknown, we made predictions for 4 possible N-terminal dimerization domain locations (Figure S2), the shortest and longest of which are shown here (Figure 6b-c). We found that the effect of N-terminal dimerization on the Bni1 polymerization rate depends on the location of the dimerization domain. The prediction histograms contain two peaks: a large peak corresponding to decreased polymerization rates and a smaller peak corresponding to a less substantial decrease (Figure S2a), no change (Figure S2b), or a mild increase (Figure S2c-d) in polymerization. Minor peaks corresponding to mild accelerations occur when the N-terminal dimerization domain is farther away from the most N-terminal binding site. This is consistent with the behavior of the model described above (Figure 5) as, for any fixed binding site location with respect to the FH2 domain (*l*_*CT*_), increasing the distance from the binding site to the N-terminal dimerization domain (*l*_*NT*_) predicted a gradual shift from deceleration to mild acceleration with N-terminal dimerization.

The best-fit parameters, as determined by maximum likelihood, fit the data, and are shown in Figure S3. Importantly, the best-fit parameters represent a regime where capture and delivery compete with one another (Figure 4d-e, pink dashed lines). Thus, when trained on experimental data, the model favored a regime where polymerization rates are sensitive to changes in both binding site accessibility and local effective concentration upon N-terminal dimerization.

### 3.5 The model can be used to predict the impact of N-terminal dimerization across multiple formin family members and hypothetical N-terminal dimerization domain placements

Using the best-fit parameters from above, we predicted the effect of N-terminal dimerization for multiple formins (Table S1). In order to assess the impact of N-terminal dimerization domain location, we made predictions for formin constructs with dimerization domains at a range of distances from the N-terminal most binding site. We note that precise locations for N-terminal dimerization domains are unknown for most formins. Therefore, we placed the dimerization domain at locations beginning at the most N-terminal binding site (*l*_*NT*_ = 0) and ending where the total FH1 domain length is 400 amino acids. We visualized these predictions as a heatmap and a family of curves (Figure 7a-b).

**Figure 7.**
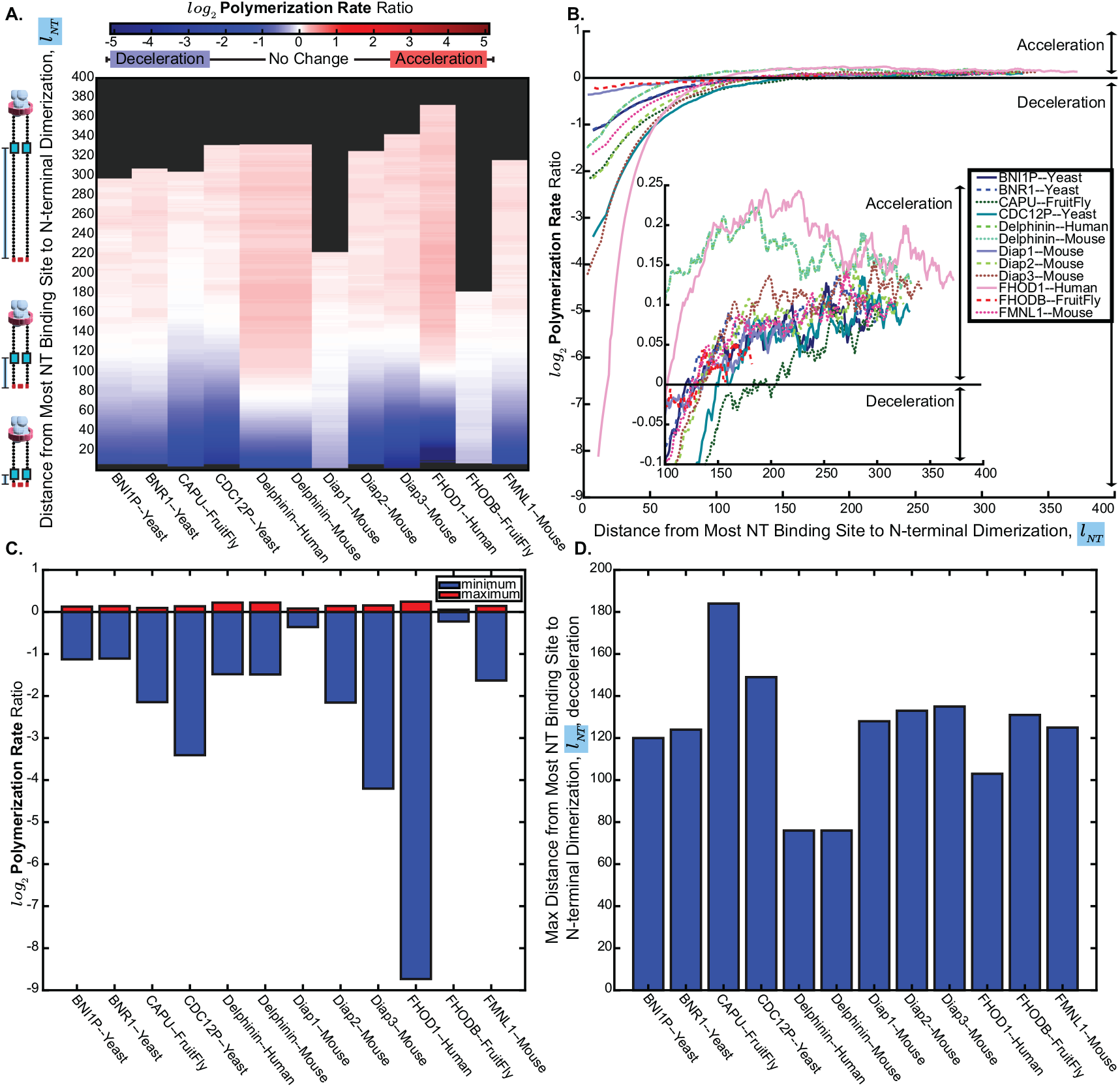
Model predicts deceleration → acceleration spectrum based on N-terminal dimerization domain placement for multiple formin family members. (a-b) Predicted polymerization rate ratios (dimerized/non-dimerized) for multiple formin family members swept across possible N-terminal dimerization domain locations, presented as either a heatmap (a) or a family of curves (b). A rolling average with a window size of 20 was taken for the ratios. Color-map values range from blue to red, indicating a deceleration or acceleration of polymerization due to N-terminal dimerization, respectively. For each formin, the N-terminal dimerization domain was placed at locations from the most N-terminal binding site (PRM, proline-rich motif) (*l*_*NT*_ = 0) and to 400 amino acids away from the FH2 domain, resulting in a smaller range of simulated values for longer formins. Simulations were run using with gating factors (*G*) listed in Table S1 and the following parameters: 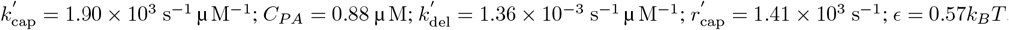. (c)The maximum (greatest acceleration, red) and minimum (greatest deceleration, blue) polymerization rate ratios (dimerized/non-dimerized) for each of the formins from the simulations in (a-b). (d) The maximum distance between the N-terminal binding site and the dimerization domain (*l*_*NT*_) at which polymerization rates only decelerate for each of the formins from the simulations in (a-b). For these measurements, rolling averages were not used, and the maximum distance was determined by finding the first point at which the polymerization rate ratio (dimerized/non-dimerized) was greater than or equal to 0.

The model predicted that as the dimerization domain moves away from the binding sites, the impact of N-terminal dimerization diminishes (Figure 7), as expected. For formin constructs with dimerization domains close to the FH1 domain, the model predicted a significant deceleration of polymerization (dark blue at the bottom of Figure 7a and bottom left of Figure 7b). For formin constructs with dimerization domains far from the FH1 domain, the model predicted essentially no change in polymerization (white and light red at the top of Figure 7a and top right of Figure 7b). The maximum predicted decelerations are much larger than the maximum predicted accelerations (Figure 7c). Such behavior is expected based on the best-fit parameters; Since the parameter regime is such that capture and delivery compete with one another, binding sites, depending on their location, can experience significant deceleration, no change, or mild acceleration (Figure 5b).

Additionally, we identified the approximate maximum dimerization domain to FH1 domain distance for which deceleration continuously occurs (Figure 7d). If dimerization domains exist within these distances from the FH1 domain, our model predicted that N-terminal dimerization will cause FH1-mediated polymerization to slow.

The predicted behavior of individual formins can be understood by inspecting the behavior of the FH1 domain simulated in the fast-timescale model (see Table S2). Formins predicted to experience decelerations of roughly an order of magnitude or more when the dimerization domain is close to the N-terminal binding site (Yeast CDC12P, Mouse Diap3, Human Fhod1; bottom left of Figure 7b) have binding sites close to the FH2 domain. These binding sites experience strong deceleration due to strong decreases in local effective concentration (bottom of Figure S4b). Formins predicted to experience decelerations less than 50% when the dimerization domain is close to the N-terminal binding site (Mouse Diap1, FruitFly FhodB; top left of Figure 7b) have N-terminal binding sites very far from the FH2 domain. These binding sites experience weaker deceleration due to the weaker decreases in local effective concentration (bottom of Figure S4c). Formins predicted to experience stronger (relative to other formins) acceleration when the dimerization domain is far from the N-terminal binding site (Human Delphilin, Mouse Delphilin, Human Fhod1; top right of Figure 7b) have binding sites close to the FH2 domain. These binding sites experience stronger (relative to other binding sites) acceleration due to more consistent mild accelerations in accessibility (top of Figure S4h).

## 4 Discussion

N-terminal dimerization in formins is poorly understood. While we have a crystal structure showing N-terminal dimerization in mDia1, our understanding of N-terminal dimerization for most formins is limited to the distant DID domain or coiled-coil predictions (7, 36). Furthermore, current literature primarily addresses the existence of N-terminal dimerization, with little insight into its impact on formin activity. Here, we found that N-terminal dimerization’s impact on FH1 domain polymer dynamics could lead to either modest speedup or dramatic slow-down of F-actin assembly. This result has important implications for our understanding of FH1-mediated actin polymerization. It suggests that experiments using non-N-terminally dimerized formins may yield results that differ from the behavior of full-length formins. Moreover, our model allows for a mechanistic explanation for the predicted changes in polymerization rates. We found that slowdowns are driven by deceleration of delivery, which occurs due to decreases in the local effective concentration of binding sites at the assembly site. Speedups are driven by acceleration of capture, which occurs due to increases in the accessibility of binding sites. The dominant effect depends on the binding site location and the overall parameter regime. The best fit to experimental data occurs in a parameter regime where capture and delivery generally compete with one another, making deceleration the most prominent effect. Using these best fit parameters to make predictions across different formins, we found that the impact of N-terminal dimerization diminishes as the dimerization domain moves farther away from the FH1 domain. Thus, we believe it unlikely that dimerization only at the DID domain, which is a considerable distance from the FH1 domain, significantly impacts polymerization rates. Knowledge of non-DID dimerization domain locations are thus crucial to making accurate predictions for the effects of N-terminal dimerization on individual formins.

The decrease in local effective concentration and increase in binding site accessibility predicted by the model were unexpected. When we use smaller, nonphysiological FH2 domain sizes, we observe our *a priori* expected behavior: increased local effective concentration (Figure 2d) and decreased binding site accessibility (Figure 3d) due to the doubling of the entropic spring force and the closer proximity of FH1 domains, respectively. However, realistic FH2 domain sizes, which are much larger than typical formin visualizations imply, change the geometry. Scientific illustrations, including those in this work, necessarily simplify and approximate biological reality. These simplifications and approximations, especially when they become the standard way of representing a specific process, impact the way we conceptualize the problem. Many schematic illustrations of formins successfully demonstrate, for example, the steps involved in the polymerization process or the domain-based architecture of a formin (5, 6, 37). However, these illustrations may not reflect accurate FH1-FH2 domain geometry, as they often show fully extended FH1 domains. For example, for a non-N-terminally dimerized formin with an FH1 domain 100 amino acids long, the FH1 domain would extend roughly 30Å (as root-mean-square distance of a gaussian chain scales with *n*^0.5^), compared to the *∼*106Å width of the FH2 domain (19, 22).

The formin superfamily is diverse and plentiful, with multiple formin genes occurring in the same species. However, each formin appears to play unique roles and have unique behavior (5, 6). For example, the 3 fission yeast formins, Cdc12, For3, and Fus1, polymerize actin with different rates and are uniquely involved in cytokinesis, cable formation, and sexual reproduction, respectively (5, 6, 38). While the reason why certain actin polymerization behavior is best suited to certain formin functions is an important open question (6), we do know that formin-mediated actin polymerization speed must be tightly controlled. This is best illustrated by the plethora of disease-causing mutations in human formins that include both gain and loss of function mutations (8). For example, focal segmental glomerulosclerosis (FSGS, scar tissue accumulation on the kidney) has been tied to both loss of function mutations in DAAM2 and gain of function mutations in INF2 (8). Increased actin polymerization is not necessarily beneficial, as evidenced by the accumulation of actin bundles seen in the FSGS-causing INF2 mutation (8). Depending on the formin and the dimerization domain placement, our results suggest that N-terminal dimerization could play a significant role in regulating the speed of actin polymerization. This could help explain the high concentration of hypertrophic cardiomyopathy-causing mutations in the putative coiled-coil domain of Fhod3 (17). Our work prompts further investigation of dimerization N-terminal to the FH1 domain: Do all formins contain an N-terminal dimerization domain? Is N-terminal dimerization constitutive? Are there other regulatory elements involved in N-terminal dimerization?

Our results also highlight the importance of the actin monomer delivery site location in understanding the mechanism by which N-terminal dimerization impacts polymerization rates. In the fast-timescale polymer model, we use a delivery site location of the FH1-FH2 domain attachment point. If a delivery site closer to the center of the FH2 domain is used, N-terminal dimerization may yield different results, most likely a dampened decrease or possible increase in local effective concentration similar to the results with smaller, nonphysiological FH2 domain widths (Figure 2a-b). Evidence for the location of the “true” physiological delivery site is lacking, but a cryo-EM structure of the formin Cdc12 with a profilin-bound binding site attached to the growing actin filament (the so-called “ring complex”) places the binding site between the attachment point and the center of the FH2 domain (35). This “off-center” delivery site supports a mechanism where N-terminal dimerization pulls FH1 density away from the delivery site, but also suggests that non-N-terminally dimerized FH1 domains, which pull FH1 density towards the attachment point, may do the same. However, this crystal structure represents the FH1 domain conformation after actin monomer addition and does not necessarily reflect the conformation of the FH1 domain prior to monomer incorporation.

A key direction for future work is to validate and test the model’s predictions against experimental data. Given the lack of well-defined posteriors from training the model on the current dataset (Figure 6), additional experimental data for model training will likely be needed to yield stronger predictions. Experimental data measuring the impacts of N-terminal dimerization both for model training and validation will thus be crucial for expanding the capabilities of our model. While experimental constructs with known N-terminal dimerization domain locations (such as those with an exogenous GST or leucine zipper) can be used for training and validation, known endogenous dimerization domain locations beyond the DID domain would enable more robust predictions for additional formins. Additionally, work towards reducing polymer dynamics simulation runtime would be ideal. One possible avenue for this is replacing the freely-jointed chain simulations with analytical mathematical expressions for local effective concentration and binding site accessibility; Obtaining mathematical expressions for polymer-dynamics problems has previously been successful (39–41).

Using simple polymer physics and an uncomplicated kinetic scheme, we developed a model of FH1-mediated polymerization that can be fit to experimental data and make predictions for multiple formins while remaining interpretable. Increasing the model’s complexity may improve fit, but would further increase the complexity of an already challenging parameter space. With the current level of complexity, we are able to pinpoint specific trends and arrive at novel mechanistic explanations. Moreover, our model could easily be expanded to explain additional poorly understood facets of formin-mediated actin polymerization, such as possible cooperation and competition between binding sites (28). Beyond formins, our model is a success story of using simple polymer physics to model and gain an understanding of IDR behavior. Future computational work may take similar approaches to modeling the hundreds of remaining IDR-containing proteins.

## Appendix A

Mean First Passage Time (MFPT) Calculations

The polymerization rates for each binding site (PRM) are computed based on the average time it takes to reach the “successful delivery” state assuming the system begins with a naked formin (free profilin-actin and unbound binding site). We compute this Mean First Passage Time (MFPT), 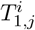, of the *i*th binding site as the sum of the residence time, 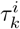, in each state preceding “successful delivery”

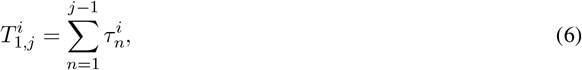

where the states are numbered sequentially with state 1 corresponding to the “naked” formin and state j to “successful delivery.”

We compute the residence times by solving a system of equations that can be simplified as

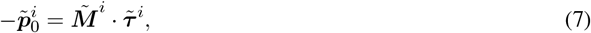

where the tilde (∼) indicates that the *j*th row and column have been dropped, 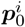 is a vector of initial conditions (1 for state 1, 0 for all other states), and ***M***^*i*^ is the transition matrix for the ith binding site.

In our model (Figure 1), “successful delivery” refers to the 3rd state.This results in the 3×3 transition matrix is

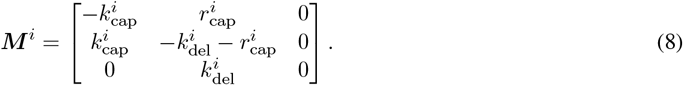

Dropping the 3rd row and column for ***M*** ^*i*^ and using Equation (7) yields

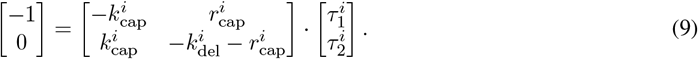

After solving for 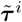, we get

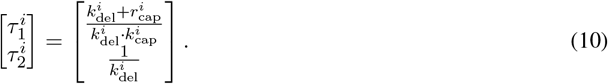

Using Equation (6), the MFPT to reach state 3 is thus:

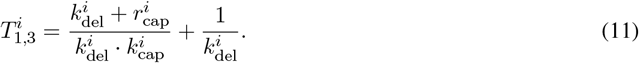

Finally, we arrive at the polymerization rate for the ith binding site by taking the reciprocal of Equation (11)

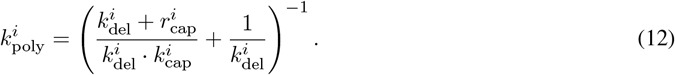

## Appendix B

**Note on** 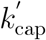 **Units**

We report 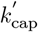 in units of based on the inclusion of *C*_PA_ in the calculation for *k*_cap_ (Equation (2)). However, we can also use a combined rate constant, 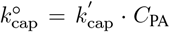, with units s^−1^. Since all model fitting was done with *C*_PA_ = 0.88 µ M^−1^, 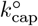 might be the more appropriate best fit parameter when translating the model to experiments with different *C*_PA_ values. In this case, 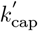 would vary based on the experimental conditions, and could be determined by taking the best fit 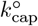 and dividing by *C*_PA_.

## 5 Data Availability

All coarse-grain polymer simulations were performed using C with analysis in MATLAB R2023a. C code available at: https://github.com/katiebogue/FH1_polymer_simulation.git DOI: 10.5281/zen-odo.17517114.

All kinetic simulations and analysis were performed in MATLAB R2023a. Code to generate Figure 4 and Figure 5 available at: https://github.com/katiebogue/formin_kinetic_model.git DOI: 10.5281/zen-odo.17517108.

## 6 Author Contributions

KB: conceptualization, data curation, formal analysis, investigation, software, visualization, writing. BC: conceptualization, writing - reviewing and editing. MQ: conceptualization, funding acquisition, supervision, writing. JA: conceptualization, funding acquisition, supervision, writing.

## 8 Acknowledgements

This work was funded by NSF DMS 2052668 to JA and NIH R01GM096133 to MQ. Computation was performed on the UC Irvine Research Cyberinfrastructure Center. Thank you to Naomi Courtemanche for Bni1 construct sequence information.

## References

[1] van der Lee, R., et al. 2014. Classification of Intrinsically Disordered Regions and Proteins. Chem. Rev. 114(13):6589–6631. doi:10.1021/cr400525m.

[2] Uversky, V. N., C. J. Oldfield, and A. K. Dunker. 2008. Intrinsically Disordered Proteins in Human Diseases: Introducing the D2 Concept. Annu. Rev. Biophys. 37:215–246. doi:10.1146/annurev.biophys.37.032807.125924.

[3] Dunker, A. K., C. J. Brown, J. D. Lawson, L. M. Iakoucheva, and Z. Obradović. 2002. Intrinsic Disorder and Protein Function. Biochemistry 41(21):6573–6582. doi:10.1021/bi012159+.

[4] Pruyne, D. 2016. Revisiting the Phylogeny of the Animal Formins: Two New Subtypes, Relationships with Multiple Wing Hairs Proteins, and a Lost Human Formin. PLOS ONE 11(10):e0164067. doi:10.1371/journal.pone.0164067.

[5] Valencia, D. A. and M. E. Quinlan. 2021. Formins. Curr. Biol. 31(10):R517–R522. doi:10.1016/j.cub.2021.02.047.

[6] Courtemanche, N. 2018. Mechanisms of formin-mediated actin assembly and dynamics. Biophys. Rev. 10(6):1553–1569. doi:10.1007/s12551-018-0468-6.

[7] Higgs, H. N. and K. J. Peterson. 2005. Phylogenetic Analysis of the Formin Homology 2 Domain. Mol. Biol. Cell 16(1):1–13. doi:10.1091/mbc.e04-07-0565.

[8] Labat-de Hoz, L. and M. A. Alonso. 2021. Formins in Human Disease. Cells 10(10):2554. doi:10.3390/cells10102554.

[9] Goode, B. L. and M. J. Eck. 2007. Mechanism and Function of Formins in the Control of Actin Assembly. Annu. Rev. Biochem. 76(1):593–627. doi:10.1146/annurev.biochem.75.103004.142647.

[10] Kovar, D. R., E. S. Harris, R. Mahaffy, H. N. Higgs, and T. D. Pollard. 2006. Control of the Assembly of ATP- and ADP-Actin by Formins and Profilin. Cell 124(2):423–435. doi:10.1016/j.cell.2005.11.038.

[11] Paul, A. and T. Pollard. 2008. The Role of the FH1 Domain and Profilin in Formin-Mediated Actin-Filament Elongation and Nucleation. Curr. Biol. 18(1):9–19. doi:10.1016/j.cub.2007.11.062.

[12] Vavylonis, D., D. R. Kovar, B. O’Shaughnessy, and T. D. Pollard. 2006. Model of Formin-Associated Actin Filament Elongation. Mol. cell 21(4):455–466. doi:10.1016/j.molcel.2006.01.016.

[13] Bryant, D., L. Clemens, and J. Allard. 2017. Computational simulation of formin-mediated actin polymerization predicts homologue-dependent mechanosensitivity. Cytoskeleton 74(1):29–39. doi:10.1002/cm.21344.

[14] Li, F. and H. N. Higgs. 2005. Dissecting Requirements for Auto-inhibition of Actin Nucleation by the Formin, mDia1. J. Biol. Chem. 280(8):6986–6992. doi:10.1074/jbc.M411605200.

[15] Higgs, H. N. 2005. Formin proteins: a domain-based approach. Trends Biochem. Sci. 30(6):342–353. doi: 10.1016/j.tibs.2005.04.014.

[16] Bor, B., C. L. Vizcarra, M. L. Phillips, and M. E. Quinlan. 2012. Autoinhibition of the formin Cappuccino in the absence of canonical autoinhibitory domains. Mol. Biol. Cell 23(19):3801–3813. doi:10.1091/mbc.e12-04-0288.

[17] Ochoa, J. P., et al. 2018. Formin Homology 2 Domain Containing 3 (FHOD3) Is a Genetic Basis for Hypertrophic Cardiomyopathy. J. Am. Coll. Cardiol. 72(20):2457–2467. doi:10.1016/j.jacc.2018.10.001.

[18] Teraoka, I. 2002. Models of Polymer Chains. John Wiley & Sons, Ltd. ISBN 978-0-471-22451-8. doi: 10.1002/0471224510.ch1.

[19] Doi, M. and S. F. Edwards. 1988. The Theory of Polymer Dynamics. International Series of Monographs on Physics. Oxford University Press, Oxford, New York. ISBN 978-0-19-852033-7.

[20] Pruyne, D., M. Evangelista, C. Yang, E. Bi, S. Zigmond, A. Bretscher, and C. Boone. 2002. Role of formins in actin assembly: nucleation and barbed-end association. Sci. (New York, N.Y.) 297(5581):612–615. doi: 10.1126/science.1072309.

[21] Courtemanche, N. and T. D. Pollard. 2012. Determinants of Formin Homology 1 (FH1) Domain Function in Actin Filament Elongation by Formins. The J. Biol. Chem. 287(10):7812–7820. doi:10.1074/jbc.M111.322958.

[22] Xu, Y., J. B. Moseley, I. Sagot, F. Poy, D. Pellman, B. L. Goode, and M. J. Eck. 2004. Crystal Structures of a Formin Homology-2 Domain Reveal a Tethered Dimer Architecture. Cell 116(5):711–723. doi:10.1016/S0092-8674(04)00210-7.

[23] Michele, C. D., P. D. L. Rios, G. Foffi, and F. Piazza. 2016. Simulation and Theory of Antibody Binding to Crowded Antigen-Covered Surfaces. PLoS Comput. Biol. 12(3):e1004752. doi:10.1371/journal.pcbi.1004752.

[24] Clemens, L., O. Dushek, and J. Allard. 2021. Intrinsic Disorder in the T Cell Receptor Creates Cooperativity and Controls ZAP70 Binding. Biophys. J. 120(2):379–392. doi:10.1016/j.bpj.2020.11.2266.

[25] Valen, D. V., M. Haataja, and R. Phillips. 2009. Biochemistry on a Leash: The Roles of Tether Length and Geometry in Signal Integration Proteins. Biophys. J. 96(4):1275–1292. doi:10.1016/j.bpj.2008.10.052.

[26] Imran, A., B. S. Moyer, A. J. Wolfe, M. S. Cosgrove, D. E. Makarov, and L. Movileanu. 2022. Interplay of Affinity and Surface Tethering in Protein Recognition. The J. Phys. Chem. Lett. 13(18):4021–4028. doi: 10.1021/acs.jpclett.2c00621.

[27] González-Foutel, N. S., et al. 2022. Conformational buffering underlies functional selection in intrinsically disordered protein regions. Nat. Struct. & Mol. Biol. 29(8):781–790. doi:10.1038/s41594-022-00811-w.

[28] Zweifel, M. E. and N. Courtemanche. 2020. Competition for delivery of profilin–actin to barbed ends limits the rate of formin-mediated actin filament elongation. J. Biol. Chem. 295(14):4513–4525. doi:10.1074/jbc.RA119.012000.

[29] Zhao, C., C. Liu, C. W. Hogue, and B. C. Low. 2014. A cooperative jack model of random coil-to-elongation transition of the FH1 domain by profilin binding explains formin motor behavior in actin polymerization. FEBS Lett. 588(14):2288–2293. doi:10.1016/j.febslet.2014.05.016.

[30] Rohatgi, A. 2022. WebPlotDigitizer. E-Mail.

[31] Eads, J. C., N. M. Mahoney, S. Vorobiev, A. R. Bresnick, K.-K. Wen, P. A. Rubenstein, B. K. Haarer, and S. C. Almo. 1998. Structure Determination and Characterization of Saccharomyces cerevisiae Profilin. Biochemistry 37(32):11171–11181. doi:10.1021/bi9720033.

[32] Metropolis, N., A. W. Rosenbluth, M. N. Rosenbluth, A. H. Teller, and E. Teller. 1953. Equation of State Calculations by Fast Computing Machines. The J. Chem. Phys. 21(6):1087–1092. doi:10.1063/1.1699114.

[33] Neal, R. M. 2011. MCMC using Hamiltonian dynamics. In Handbook of Markov Chain Monte Carlo. Chapman & Hall\CRC. doi:10.1201/b10905.

[34] Press, W. H. 2007. Numerical Recipes 3rd Edition: The Art of Scientific Computing. Cambridge University Press. ISBN 978-0-521-88068-8.

[35] Oosterheert, W., M. Boiero Sanders, J. Funk, D. Prumbaum, S. Raunser, and P. Bieling. 2024. Molecular mechanism of actin filament elongation by formins. Science 384(6692):eadn9560. doi:10.1126/science.adn9560.

[36] Nezami, A., F. Poy, A. Toms, W. Zheng, and M. J. Eck. 2010. Crystal Structure of a Complex between Amino and Carboxy Terminal Fragments of mDia1: Insights into Autoinhibition of Diaphanous-Related Formins. PLOS ONE 5(9):e12992. doi:10.1371/journal.pone.0012992.

[37] Paul, A. S. and T. D. Pollard. 2009. Energetic Requirements for Processive Elongation of Actin Filaments by FH1FH2-formins. The J. Biol. Chem. 284(18):12533–12540. doi:10.1074/jbc.M808587200.

[38] Scott, B. J., E. M. Neidt, and D. R. Kovar. 2011. The functionally distinct fission yeast formins have specific actin-assembly properties. Mol. Biol. Cell 22(20):3826–3839. doi:10.1091/mbc.E11-06-0492.

[39] Bell, S. and E. M. Terentjev. 2017. Kinetics of Tethered Ligands Binding to a Surface Receptor. Macromolecules 50(21):8810–8815. doi:10.1021/acs.macromol.7b01742.

[40] Shimamura, M. K. and T. Deguchi. 2002. Knot complexity and the probability of random knotting. Phys. Rev. E 66(4):040801. doi:10.1103/PhysRevE.66.040801.

[41] Segall, D. E. 2006. Volume-Exclusion Effects in Tethered-Particle Experiments: Bead Size Matters. Phys. Rev. Lett. 96(8). doi:10.1103/PhysRevLett.96.088306.

[42] Shimada, A., M. Nyitrai, I. R. Vetter, D. Kühlmann, B. Bugyi, S. Narumiya, M. A. Geeves, and A. Wittinghofer. 2004. The Core FH2 Domain of Diaphanous-Related Formins Is an Elongated Actin Binding Protein that Inhibits Polymerization. Mol. Cell 13(4):511–522. doi:10.1016/S1097-2765(04)00059-0.

[43] Harris, E. S., I. Rouiller, D. Hanein, and H. N. Higgs. 2006. Mechanistic Differences in Actin Bundling Activity of Two Mammalian Formins, FRL1 and mDia2*. J. Biol. Chem. 281(20):14383–14392. doi:10.1074/jbc.M510923200.

[44] Schönichen, A., et al. 2013. FHOD1 is a combined actin filament capping and bundling factor that selectively associates with actin arcs and stress fibers. J. Cell Sci. 126(8):1891–1901. doi:10.1242/jcs.126706.

[45] Moseley, J. B. and B. L. Goode. 2005. Differential Activities and Regulation of Saccharomyces cerevisiae Formin Proteins Bni1 and Bnr1 by Bud6 *. J. Biol. Chem. 280(30):28023–28033. doi:10.1074/jbc.M503094200.

[46] Bremer, K. V., C. Wu, A. A. Patel, K. L. He, A. M. Grunfeld, G. F. Chanfreau, and M. E. Quinlan. 2024. Formin tails act as a switch, inhibiting or enhancing processive actin elongation. J. Biol. Chem. 300(1):105557. doi:10.1016/j.jbc.2023.105557.

[47] Silkworth, W. T., K. L. Kunes, G. C. Nickel, M. L. Phillips, M. E. Quinlan, and C. L. Vizcarra. 2018. The neuron-specific formin Delphilin nucleates nonmuscle actin but does not enhance elongation. Mol. Biol. Cell 29(5):610–621. doi:10.1091/mbc.E17-06-0363.

