## Supplemental Material for "Impact of N-terminal dimerization on formin homology 1 domain polymer dynamics and actin assembly"

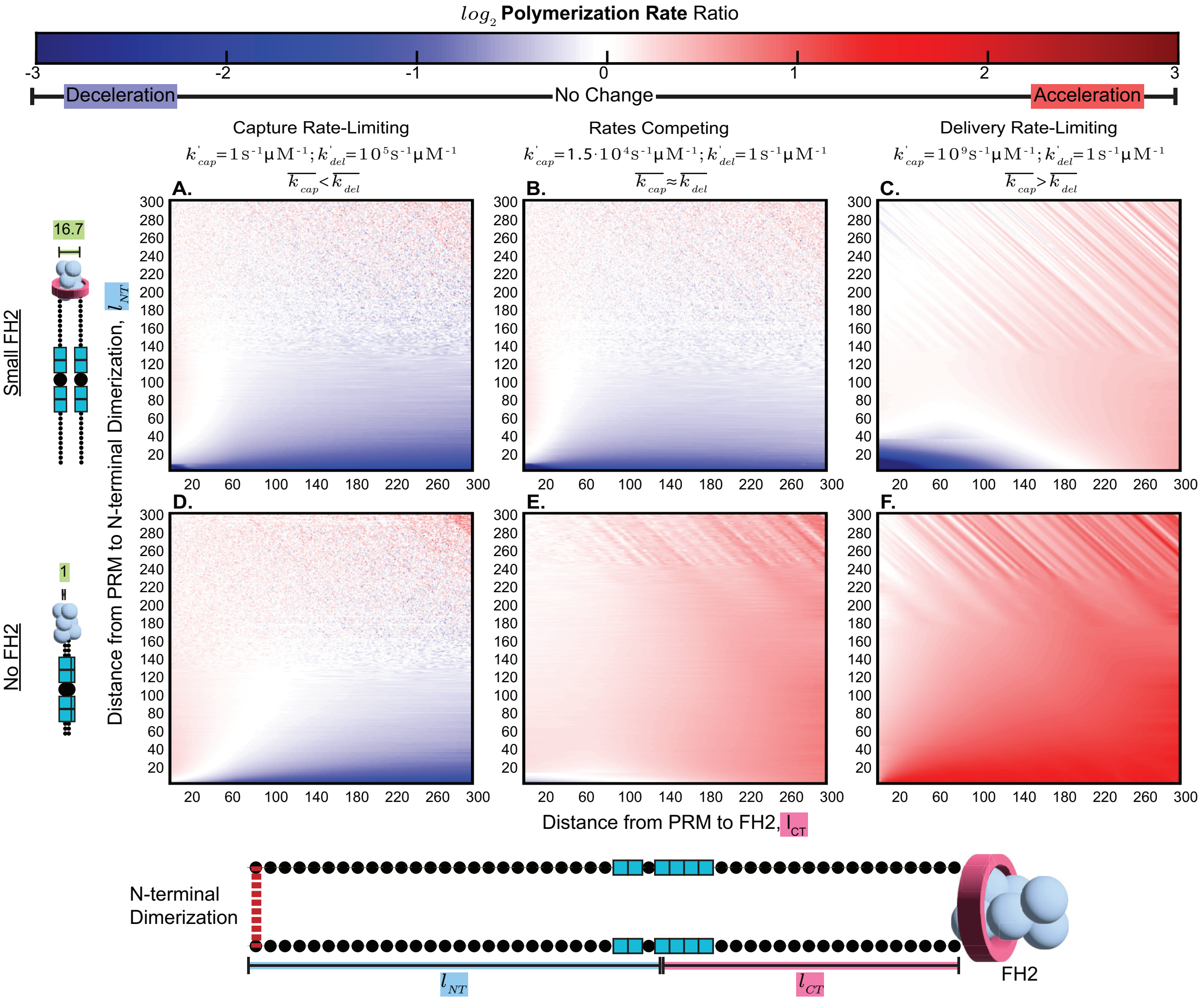


Figure S1: **N-terminal dimerization can result in acceleration or deceleration of polymerization based on parameter regime.** Combining the polymer behavior model (Figure 2 and Figure 3) with the kinetic model (Figure 4), we can predict the impact of N-terminal dimerization on overall polymerization rates. The ratio (dimerized/non-dimerized) of polymerization rates for formins with a single binding site (PRM, proline-rich motif) are plotted as heatmaps with respect to the location of the binding site: distance from binding site to FH2 domain ($l_{CT}$, x-axis) and distance from binding site to N-terminal dimerization site ($l_{NT}$, y-axis). Color-map values range from blue to red, indicating a deceleration or acceleration of polymerization due to N-terminal dimerization, respectively. All regimes used the following parameters: $r^i_{\text{cap}} = 3.35\ \text{s}^{-1}$; $G = 1$; $C_{PA} = 1$ $\mu$M. $k^{'}_{\text{cap}}$ and $k^{'}_{\text{del}}$ values were selected to showcase the behavior when capture is rate-limiting (a, d; $k^{'}_{\text{cap}} = 1\ \text{s}^{-1}\,\mu\text{M}^{-1}$, $k^{'}_{\text{del}} = 10^5\ \text{s}^{-1}\,\mu\text{M}^{-1}$), when delivery is rate-limiting (c, f;, $k^{'}_{\text{cap}} = 1.5 * 10^4\ \text{s}^{-1}\,\mu\text{M}^{-1}$, $k^{'}_{\text{del}} = 1\ \text{s}^{-1}\,\mu\text{M}^{-1}$), and when capture and delivery are competing (b, e;, $k^{'}_{\text{cap}} = 10^9\ \text{s}^{-1}\,\mu\text{M}^{-1}$, $k^{'}_{\text{del}} = 1\ \text{s}^{-1}\,\mu\text{M}^{-1}$). Simulations were run for 2 different FH2 domain sizes: (a-c) 16.7 amino acids and (d-f) 1 amino acid.

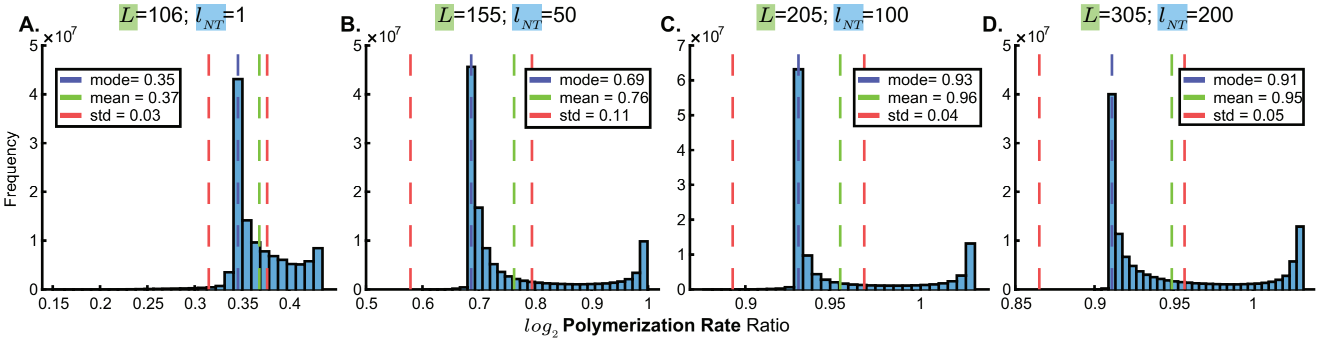


Figure S2: **Predicting the effect of N-terminal dimerization on Bni1 using MCMC posteriors.** Predicted polymerization rate ratios (dimerized/non-dimerized) for Bni1 using parameters from the posterior distribution of the MCMC fit to single binding site (PRM, proline-rich motif) Bni1 data. Simulations were run for 5 different binding site locations ($l_{NT}$) and FH1 domain lengths ($L$): (a) $L = 106$, $l_{NT} = 1$, (b) $L = 155$, $l_{NT} = 50$, (c) $L = 205$, $l_{NT} = 100$, (d) $L = 305$, $l_{NT} = 200$, The histogram mode (blue dashed line), mean (green dashed line), and 1 standard deviation from the mode (red dashed lines) are plotted on the histograms.

Table S1: Formins used in N-terminal dimerization predictions

| Formin | UniProt ID | Gating Factor | Ref. | Number of binding sites (PRMs) | Mean binding site (PRM) Size | Number of Prolines | FH1 domain Length |
|---|---|---|---|---|---|---|---|
| Diap1–Mouse | O08808 | 1 | (28) | 14 | 6.571429 | 92 | 181 |
| Diap2–Mouse | O70566 | 0.2 | (28) | 5 | 8.8 | 44 | 82 |
| Diap3–Mouse | Q9Z207 | 0.26 | (42) | 3 | 9 | 27 | 61 |
| CAPU–FruitFly | Q24120 | 1 | (28) | 5 | 9.2 | 46 | 101 |
| FMNL1–Mouse | Q9JL26 | 0.55 | (43) | 4 | 9.25 | 37 | 91 |
| FHOD1–Human | Q9Y613 | 0.04 | (44) | 2 | 9 | 18 | 34 |
| BNR1–Yeast | P40450 | 1 | (45) | 4 | 8.25 | 33 | 100 |
| CDC12P–Yeast | Q10059 | 0.05 | (28) | 2 | 8.5 | 17 | 76 |
| BNI1P–Yeast | P41832 | 0.5 | (28) | 4 | 9.5 | 38 | 110 |
| FHODB–FruitFly | Q9VSQ0 | 0.5 | (46) | 2 | 10.5 | 21 | 226 |
| Delphilin–Human | A4D2P6 | 0.04 | (47) | 4 | 6.5 | 26 | 71 |
| Delphilin–Mouse | Q0QWG9 | 0.03 | (47) | 4 | 6.5 | 26 | 71 |

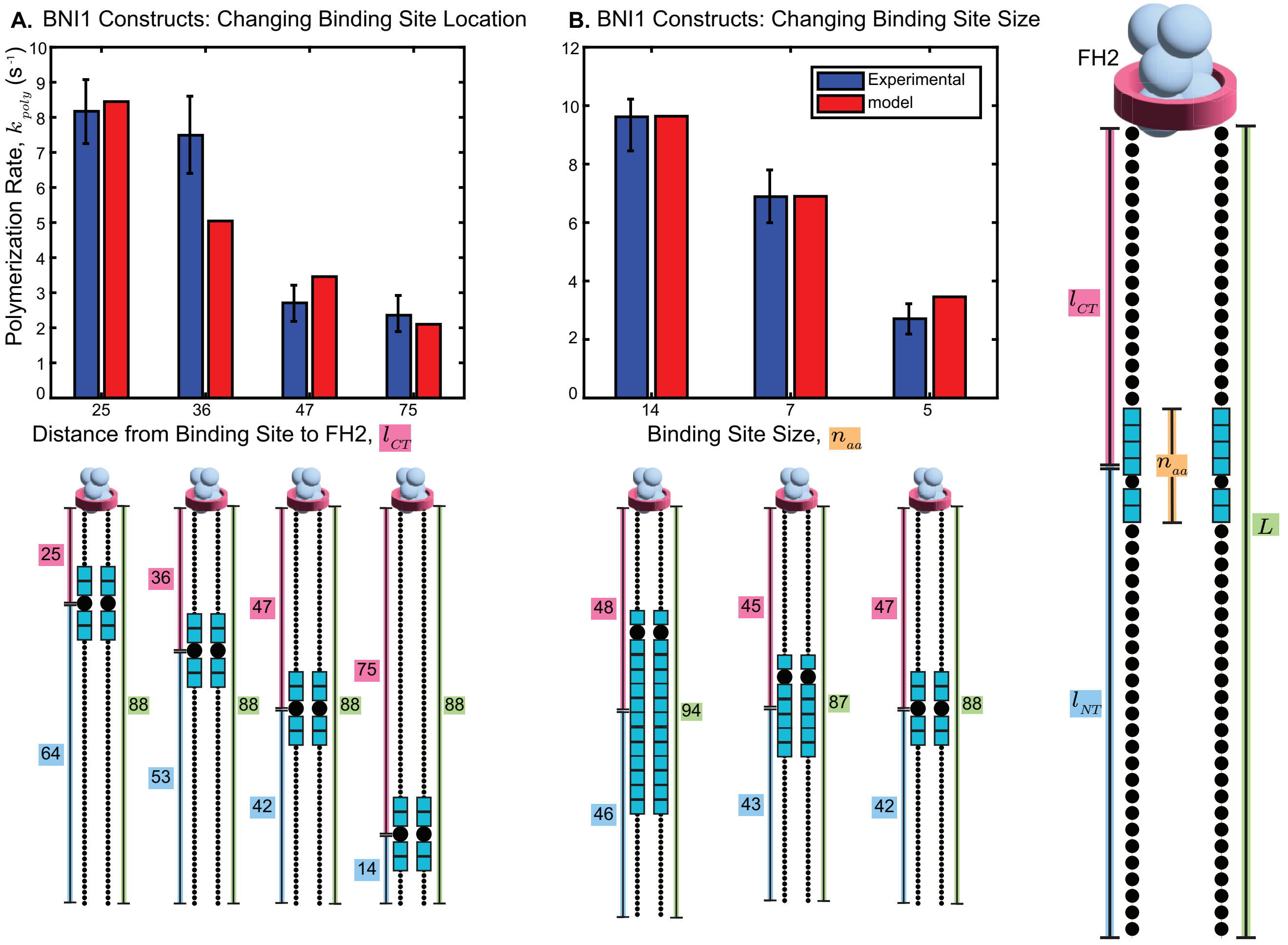


Figure S3: **Fit to experimental single binding site Bni1 data using best-fit parameters from MCMC fit.** Experimental (blue) and simulated (red) polymerization rates for single binding site (PRM, proline-rich motif) Bni1 constructs shown as side-by-side bar graphs. Constructs include those with the same binding site (PPAPP) at different locations along the FH1 domain (a) and those with different binding sites all at (roughly) the same location along the FH1 domain (b). Experimental values and error bars are taken from figures 3 and 4 of Courtemanche and Pollard (21) at profilin concentrations of 5μM. Simulations were run using the best-fit parameters from the posterior distribution of the MCMC (Figure 6): $k^{'}_{\text{cap}} = 1.90 \times 10^3$ $\text{s}^{-1}\,\mu\text{M}^{-1}$; $C_{PA} = 0.88\,\mu\text{M}$; $k^{'}_{\text{del}} = 1.36 \times 10^{-3}$ $\text{s}^{-1}\,\mu\text{M}^{-1}$; $r^{'}_{\text{cap}} = 1.41 \times 10^3$ $\text{s}^{-1}$; $\epsilon = 0.57k_BT$; $G = 0.5$. The best-fit parameters result in a RMSE of 0.639 subunits/s (compared to the experimental average SEM of 0.761 subunits/s) for the data changing binding site location (a) and a RMSE of 0.250 subunits/s (compared to the experimental average SEM of 0.769 subunits/s) for the data changing binding site site (b). Note that none of these constructs are N-terminally dimerized.

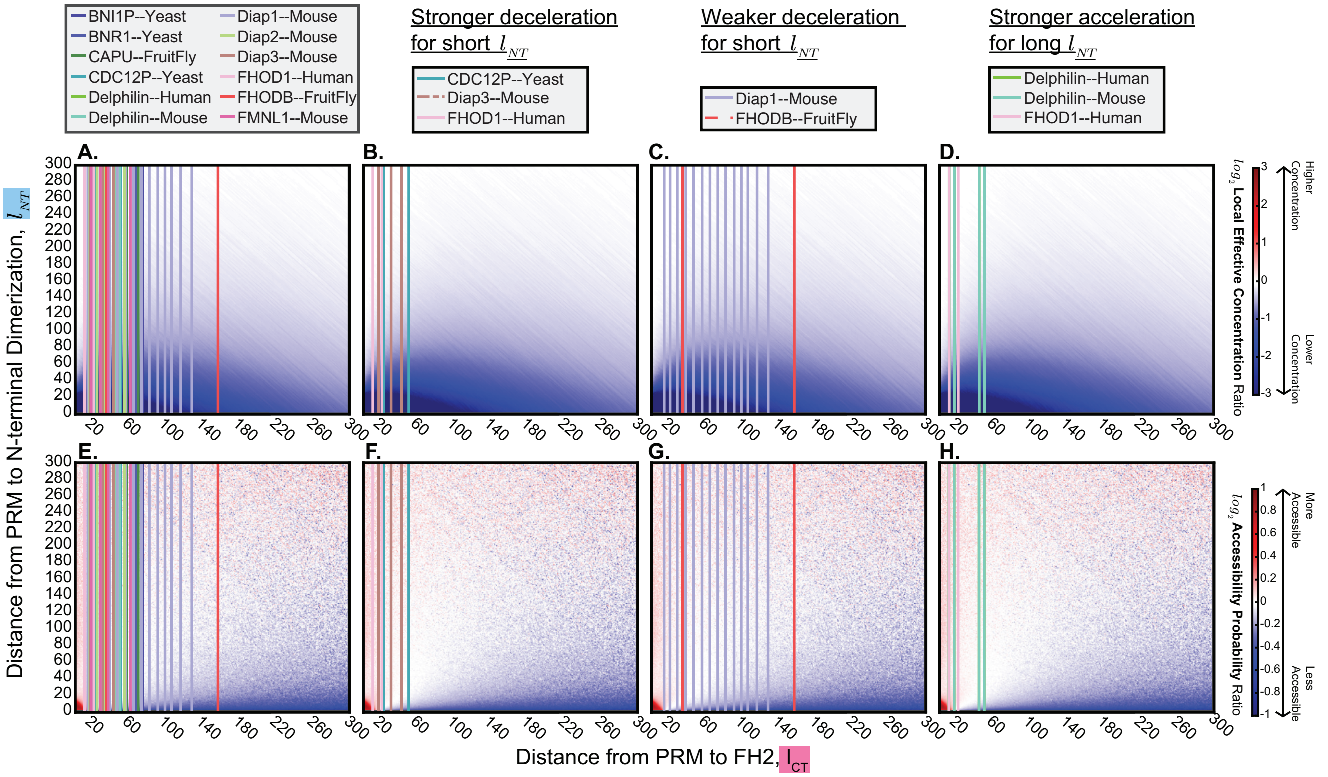


Figure S4: **Visualizing polymer behavior for binding sites based on the predicted effects of N-terminal dimerization.** The location of each binding site (PRM, proline-rich motif) for each formin (a,e) used in predictions (Figure 7) is plotted over the heatmap of ratios (dimerized/non-dimerized) of polymer behavior parameters (Figure 2c, Figure 3c) with an FH2 domain size of 35.5 amino acids (from crystal structure of the Bni1 FH2 domain dimer, Xu et al. (22), PDB: 1UX5). Since the N-terminal dimerization domain locations for each formin are unknown, the binding site locations are plotted as vertical lines at the distance from the FH2 domain to the binding site ($l_{CT}$). (a-d) Simulated local effective concentration of binding sites at the growing end ($P_{\mathrm{r}}^{i,0}$). Higher (red) or lower (blue) local concentration of binding sites at the growing end enhances or impairs the rate of delivery, respectively. (e-h) Simulated binding site accessibility ($1 - P_{\mathrm{occ}}^{i}$). Increased (red) or decreased (blue) accessibility of binding sites enhances or impairs the rate of capture, respectively. Formins with extreme predicted effects of N-terminal dimerization are grouped by behavior and shown in additional plots: (b, f) formins that are predicted to experience greater deceleration when the N-terminal binding site is near the N-terminal dimerization domain (bottom left of Figure 7b; Yeast CDC12P, Mouse Diap3, Human Fhod1); (c, g) formins that are predicted to experience less deceleration when the N-terminal binding site is near the N-terminal dimerization domain (top left of Figure 7b; Mouse Diap1, Fruit Fly FhodB); (d, h) formins that are predicted to experience greater acceleration when the N-terminal binding site is far from the N-terminal dimerization domain (top right of Figure 7b; Human Delphilin, Mouse Delphilin, Human Fhod1).

Table S2: Polymer behavior of binding sites based on the predicted effects of N-terminal dimerization

| Formins | FH1 Domain Sequence Feature | Predicted Binding Site Behavior | Predicted Polymerization Behavior |
|---|---|---|---|
| Yeast CDC12P, Mouse Diap3, Human Fhod1 | Binding sites are close to the FH2 domain | Strong decreases in local effective concentration (bottom of Figure S4b) | Strong deceleration when the dimerization domain is close to the N-terminal binding site (bottom left of Figure 7b) |
| Mouse Diap1, FruitFly FhodB | N-terminal binding sites are very far from the FH2 domain | Weaker decreases in local effective concentration (bottom of Figure S4c) | Weak deceleration when the dimerization domain is close to the N-terminal binding site (top left of Figure 7b) |
| Human Delphilin, Mouse Delphilin, Human Fhod1 | Binding sites are close to the FH2 domain | Consistent mild accelerations in accessibility (top of Figure S4h) | Stronger acceleration when the dimerization domain is far from the N-terminal binding site (top right of Figure 7b) |

Table S3: Symbols

| Rates | | |
|---|---|---|
| Symbol | Description | Units |
| $k_{\text{poly}}$ | polymerization rate for the full formin | $\text{s}^{-1}$ |
| $k^i_{\text{poly}}$ | polymerization rate for the binding site (PRM) located i amino acids away from the FH2 domain | $\text{s}^{-1}\,(\text{PRM})^{-1}$ |
| $k_{\text{cap}}$ | capture rate for the full formin | $\text{s}^{-1}$ |
| $k^i_{\text{cap}}$ | capture rate for the binding site (PRM) located i amino acids away from the FH2 domain | $\text{s}^{-1}\,(\text{PRM})^{-1}$ |
| $k_{\text{del}}$ | delivery rate for the full formin | $\text{s}^{-1}$ |
| $k^i_{\text{del}}$ | delivery rate for the binding site (PRM) located i amino acids away from the FH2 domain | $\text{s}^{-1}\,(\text{PRM})^{-1}$ |
| $r_{\text{cap}}$ | reverse capture rate for the full formin | $\text{s}^{-1}$ |
| $r^i_{\text{cap}}$ | reverse capture rate for the binding site (PRM) located i amino acids away from the FH2 domain | $\text{s}^{-1}\,(\text{PRM})^{-1}$ |

| Rate Constants | | |
|---|---|---|
| Symbol | Description | Units |
| $k'_{\text{cap}}$ | capture rate constant | $\text{s}^{-1}\,\mu\text{M}^{-1}$ |
| $k'_{\text{del}}$ | delivery rate constant | $\text{s}^{-1}\,\mu\text{M}^{-1}$ |
| $r'_{\text{cap}}$ | reverse capture rate constant | $\text{s}^{-1}$ |
| $\epsilon$ | reverse capture exponential scaling constant | n/a |

| Rate-Modifying Variables | | |
|---|---|---|
| Symbol | Description | Units |
| $P^i_{occ}$ | occlusion probability at the binding site (PRM) located i amino acids away from the FH2 domain | n/a |
| $P^0_{occ}$ | occlusion probability at the FH2/barbed end | n/a |
| $1 - P^i_{occ}$ | accessibility probability at the binding site (PRM) located i amino acids away from the FH2 domain | n/a |
| $1 - P^0_{occ}$ | accessibility probability at the FH2/barbed end | n/a |
| $P^{a,b}_r$ | probability density of location a at location b | $\mu\text{M}$ |
| $P^{i,0}_r$ | probability density of the binding site (PRM) located i amino acids away from the FH2 domain at the FH2 domain | $\mu\text{M}$ |
| $N$ | number of rods in a polymer simulation | n/a |
| $C_{\text{PA}}$ | concentration of profilin-actin in solution | $\mu\text{M}$ |
| $G$ | gating factor (0-1) | n/a |

| Length and Distance Notation | | |
|---|---|---|
| Symbol | Description | Units |
| $n_{aa}$ | total number of amino acids in a binding site (PRM) | amino acids |
| $n_p$ | number of prolines in a binding site (PRM) | amino acids |
| $L$ | length (number of amino acids) of a formin from FH2 domain to NTDd | amino acids |
| $l_{NT}$ | length (number of amino acids) from a binding site (PRM) to NTDd | amino acids |
| $l_{CT}$ | length (number of amino acids) from a binding site (PRM) to FH2 domain | amino acids |
| $L_k$ | Kuhn length | n/a |

Table S4: Terms

| Term | Description |
|---|---|
| polymerization | actin monomer addition to an actin filament (not including nucleation) |
| capture | binding site (PRM) binding to profilin-actin |
| delivery | binding site (PRM)-profilin-actin reaching the FH2 domain (loop closure) |
| polymer simulation | simulation of a random walk that produces polymer statistics; does not include formin-specific information |
| polymer statistics | the results of a polymer simulation |
| kinetic simulation | simulation of FH1-mediated polymerization, applying polymer statistics to a specific formin/construct |
| polymer model | a set of parameters, equations, set-up, etc for running polymer simulations |
| kinetic model | the set of steps (and thus equations) and their parameters used to calculate rates in a kinetic simulation |
| filament model | the method by which binding site (PRM) locations, binding site (PRM) sizes, FH1 domain N-termini, and FH1 domain C-termini are defined for a formin construct) |
| model | refers to the combination of kinetic, polymer, and filament models used to model FH1-mediated actin polymerization |
| NTD | N-terminal dimerization, the process |
| NTDd | N-terminal dimerization domain, the location |
| binding site (PRM) | proline-rich motif, the profilin-actin binding sites along the FH1 domain |
| occlusion probability | the probability that the simulated polymer conformation is such that the specified location is occluded (unavailable for binding) |
| accessibility probability | the probability that the simulated polymer conformation is such that the specified location is accessible (available for binding) |
| probability density | the probability that the simulated polymer conformation is such that one location is within a specified radius of another location (usually that a binding site is at FH2); proxy for local concentration |
| gating factor | the experimentally determined ratio of polymerization rates of FH2-only formin-mediated polymerization to actin alone; here, the probability that the FH2 domain is in a conformation that allows delivery |
| Kuhn length | the length of each rod in a freely-jointed chain |
| delivery location | the coordinates of the attachment point of the FH1 domain and the FH2 domain used when computing $P_r^{i,0}$ in polymer simulations |
| non-dimerized | refers to rates, parameters, simulations, etc where the FH1 domain is dimerized at the FH2 domain but not the NTDd |
| dimerized | refers to rates, parameters, simulations, etc where the FH1 domain is dimerized at both the FH2 domain and the NTDd |
| interruption | non-proline amino acid(s) within binding sites (PRMs) |
